# Critical contribution of mitochondria in the development of cardiomyopathy linked to desmin mutation

**DOI:** 10.1101/2023.09.14.557734

**Authors:** Yeranuhi Hovhannisyan, Zhenlin Li, Domitille Callon, Rodolphe Suspène, Vivien Batoumeni, Alexis Canette, Jocelyne Blanc, Hakim Hocini, Cécile Lefebvre, Nora El-Jahrani, Aurore L’honoré, Ekaterini Kordeli, Paul Fornes, Jean-Paul Concordet, Gérard Tachdjian, Anne-Marie Rodriguez, Jean-Pierre Vartanian, Anthony Béhin, Karim Wahbi, Pierre Joanne, Onnik Agbulut

## Abstract

Beyond the observed alterations in cellular structure and mitochondria, the cellular mechanisms linking genetic mutations to the development of heart failure in patients affected by desmin defects remain unclear due, in part, to the lack of relevant human cardiomyocyte models. We investigated the role of mitochondria using cardiomyocytes derived from human induced pluripotent stem cells carrying the heterozygous *DES*^E439K^ desmin mutation, that were either isolated from a patient or generated by gene editing. To increase physiological relevance, cells were either cultured on an anisotropic surface to obtain elongated and aligned cardiomyocytes, or as spheroids to create a micro- tissue. When applicable, results were confirmed with heart biopsies from the family harboring *DES*^E439K^ mutation. We show that mutant cardiomyocytes reproduce critical defects in mitochondrial architecture, respiratory capacity and metabolic activity as observed in patient’s heart tissue. To challenge the pathological mechanism, normal mitochondria were transferred inside the mutant cardiomyocytes. This treatment restored mitochondrial and contractile functions. This work demonstrates the crucial role of mitochondrial abnormalities in the pathophysiology of desmin-related cardiomyopathy, and opens-up new potential therapeutic perspectives.

## Introduction

Desmin-related myofibrillar myopathy (MFM1, OMIM:601419) represents a group of skeletal and cardiac muscle disorders caused by mutations in the desmin-encoding *DES* gene. Desmin is the major component of intermediate filaments (IFs) in striated and smooth muscle cells and tissues and plays an essential role in the tensile strength and integrity of muscle fibers^1–4^. In humans, about a hundred mutations of the *DES* gene have been identified which usually disrupt the ability of desmin to form filamentous networks and/or to interact with partner proteins^3,5^. Consequently, desmin mutations alter the mechanical properties of the desmin network, resulting in multiple functional and structural abnormalities in muscle cells, leading to progressive skeletal myopathy and cardiomyopathy, the most common clinical manifestations of MFM1^6^. Impairment of desmin network is also closely associated to the etiology of many striated muscle pathologies. It is worth to note that desmin IFs remodeling also occurs when proteostasis is disturbed, during inflammatory process, or during aging^7–9^.

Beyond its role in muscle cell integrity and organization, several studies conducted with genetically modified cell lines carrying human desmin mutations or in desmin knockout mice indicated that desmin critically modulates mitochondria functions^10–15^. Importantly, it has been noticed that mutated desmin leads to severe mitochondria abnormalities that directly contribute to MFM1 development. Consistent with this finding, abnormalities in the distribution and morphology of mitochondria as well as in their respiratory function have been reported with *DES* mutations using transiently transfected human or animal cells carrying human *DES* mutations^15,16^, transgenic mouse hearts with *DES* mutations^17^ as well as human heart and skeletal biopsy samples of MFM1 patients^18–20^. Moreover, several publications describe that *DES* mutations are associated with Cytochrome c oxidase (COX)- negative fibers and decreased activity of mitochondrial respiratory chain enzymes^19,20^. Whether and how desmin regulates mitochondria network and function is far from being understood. Several reports indicated that desmin could mediate this effect through its physical interaction with mitochondria. Indeed, desmin can interact with mitochondria through either cytolinker proteins such as plectin isoform 1b^21–23^ or voltage-dependent anion channel (VDAC)^24,25^, or directly through its N-terminal domain^26^. Despite these previous studies pointing out that mitochondria are key players in the development of MFM1, none of them have used patient-derived cardiomyocytes or human cardiac organoids, at the best patient muscle biopsies or transgenic mice or transitory transfected cells carrying *DES* mutations have been used up to now. Therefore, knowledge in this field has been hampered by the lack of a suitable human cardiomyocyte experimental model to investigate the heart disease of MFM1.

To circumvent this limitation and address whether mitochondria is a critical target of desmin mutation in cardiomyocytes from MFM1 patients, we generated cardiomyocytes from induced pluripotent stem cells (iPSC) of a patient carrying heterozygous *DES*^E439K^ mutation. Using 2D and 3D models, we examined the structural and functional impairments of cardiomyocytes generated from *DES*^E439K^ patient iPSC (E439K-CMs) and compared to cardiomyocytes generated from healthy donors (Control- CMs) as well as from an isogenic pair carrying the same mutation introduced in healthy iPSC by CRISPR/Cas9 (Iso-E439K-CMs). We then focused on investigation of mitochondrial abnormalities by oxygraphy measurement as well as immunolocalization, western-blot and electron microscopy analyses. Finally, we challenged the impact of mitochondrial abnormalities as a main mechanism in the emergence of desmin-related cardiomyopathy in *DES*^E439K^ patients by implementing a treatment with extracellular vesicles (EVs) containing healthy mitochondria. Here we show that this treatment was able to significantly restore mitochondrial respiration and increase contractility of E439K-CMs. In summary, our study highlights the deleterious effect of *DES*^E439K^ and the crucial role of mitochondrial abnormalities in pathophysiology of MFM1.

## Methods

The study was conducted with samples from 2 patients (CII and CIII), from a French family with several members suffering myofibrillar myopathy (MFM1) (see **Figure-1**). Affected patients present a skeletal myopathy associated with arrhythmogenic DCM with complete atrioventricular block and atrial fibrillation. They have been diagnosed with a heterozygous pathogenic variant in the *DES* gene, namely a G-to-A transition at c.1315 (codon 439) in exon 8, which resulted in a substitution of glutamic acid by lysine in position 439 (*DES*^E439K^) at the C-terminal end of desmin. Such replacement of a negatively charged amino acid by a positively charged one is considered as a pathological modification that can affect desmin structure and its interactome^27,28^.

**Figure 1.**
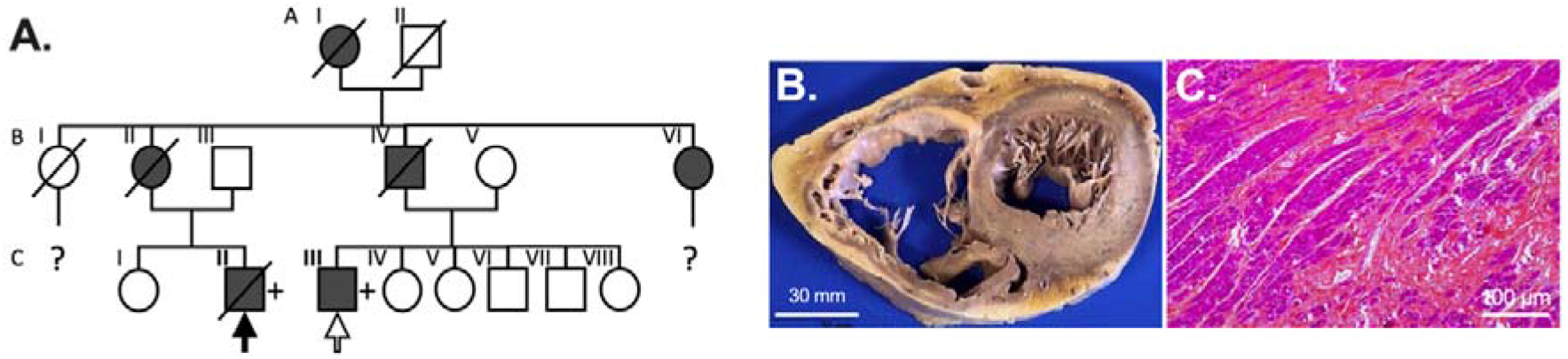
Familial myofibrillar myopathy with heterozygous *DES*^E439K^ variant. **(A)** Pedigree of the affected family. Circles and squares represent female and male subjects, respectively. Solid symbols show patients with myopathy and cardiomyopathy. Crossed-out symbols stand for deceased subjects. The + symbol represents patient with presence of the pathogenic heterozygous DES variant. Solid arrow indicates the family member whose cardiac biopsies were used for the histological and biochemical analysis. Empty arrow indicates the family member whose peripheral blood mononuclear cells were used to generate iPSC clones. **(B)** Biventricular transversal heart slice of cardiac samples of the Index case CII (solid arrow in A) showing dilation of both ventricles. **(C)** Hematoxylin-eosin safran staining of formalin fixed left ventricle section showing extensive fibrosis.

### Human tissue samples

In addition to post-mortem heart samples of the index case CII (see **Figure-1A**) carrying a heterozygous c.1315G>A (E439K) desmin mutation^29^, post-mortem heart samples from five control healthy patients were obtained after anonymization by the Platform Center of Biological Resources of the Department of Pathology (Academic Hospital of Reims, France). For each patient, authorizations to perform autopsy and sampling were obtained from the French Biomedicine Agency. Blood samples from patient CIII were collected by Phenocell SAS (Grasse, France) and peripheral blood mononuclear cells (PBMC) were freshly collected using the SepMate (Stemcell Technologies) protocol. The generation of human iPSCs were performed with approval from the French Research Ministry in Phenocell SAS (agreement number: #AC-2013-1973). Formal informed consent was obtained from the different patients included in this study.

### Generation and culture of iPSCs

PBMC of the patient CIII were reprogrammed into iPSCs using a non-integrative method (Epi5™ Episomal iPSC Reprogramming Kit, Thermo Fisher Scientific). Two clones (c2 and c15) were isolated and used in this study. Gibco Human Episomal iPSC clone (Control 1, Thermo Fisher Scientific) and CW30318 (Control 2, Fuji Cellular Dynamics) derived from healthy individuals without any cardiac pathology, were used as controls. All iPSC clones were cultured and amplified on Matrigel-coated (Corning Life Sciences) culture dishes with daily replacement of mTeSR1 (Stemcell Technologies) and passaged (1:30 ratio) when approximately 80% confluence using ReLeSR (Stemcell Technologies).

### Generation of an isogenic pair

Control 1 iPSC cultured with mTesR1 was detached using Accutase® (Stemcell Technologies) and electroporated using the Neon® electroporation system (Thermo Fisher Scientific) with 3 different plasmids diluted in resuspension RB_buffer: (1) 1000 ng/µL of pCMV_AncBE4max_P2A_GFP^30^, encoding a nucleus-targeted chimeric protein composed of a cytidine deaminase and a Cas9 nickase, 2x500 ng/µL of pBlueScript-U6sgRNA^31^ encoding single guide RNA specific for *DES* or encoding single guide RNA specific for *ATP1A1* (used for a positive selection of transformed clones)^32^. The sequence of guide RNAs for *DES* to generate E439K mutation was 5’-CTcAGAACCCCTTTGCTC-3’ and the sequence of guide RNAs for *ATP1A1* to generate point mutation to induce resistance to ouabain was 5’-cATCCAAGCTGCTACAGAAGG-3’^33^. Electroporation was performed at a voltage of 1300 V with 2 pulses on 150 000 cells resuspended in resuspension RBuffer for a 10 µl neon tip. After electroporation, the cells were transferred into pre-warmed mTeSR1 medium on Matrigel-coated plates, cultured for 3 days at which 1 µg/ml ouabain (Enzo Life Sciences) was added for clonal selection during 3 more days. The resistant cells were sequenced to confirm or deny the success of genomic editing. Single cell cloning was then performed using CloneR (Stem Cell Technologies) following the manufacturer’s instructions. pCMV_AncBE4max_P2A_GFP was a gift from David Liu (Addgene plasmid #112100; http://n2t.net/addgene:112100; RRID:Addgene_112100) and pBluescript-U6sgRNA empty was a gift from Eugene Yeo (Addgene plasmid #74707; http://n2t.net/addgene:74707; RRID:Addgene_74707).

### Generation of iPSC-derived human cardiomyocytes (iPSC-CM)

Cardiac differentiation (**Figure-2A**) is based on the GiWi protocol published in 2012^34^. At 80% confluence, iPSCs were detached with Accutase®, counted and then seeded at optimal density. Two days later, cardiac differentiation was initiated by replacing mTeSR1 with RPMI medium (Thermo Fisher Scientific) enriched with 2% insulin-free B27 supplement (Thermo Fisher Scientific) containing 9 µM CHIR99021 (Selleckchem). This medium was replaced with RPMI supplemented insulin-free B27 after 24 hours. Two days later, cells were treated with RPMI enriched with insulin-free B27 and 5 µM IWP2 (Tocris Bioscience) for 48 hours and then the medium was replaced with RPMI enriched with insulin-free B27. Two days later, the culture medium was replaced by RPMI enriched with 2% B27 supplement and was changed every two days. From day 12, 1μM dexamethasone (Merck) and 100nM triiodothyronine hormone (Merck) were added to the medium (maturation medium). At days 19-20 after initial detachment, cardiomyocytes were detached with TrypLE Select 1X (Thermo Fisher Scientific) (incubation at 37°C for 10 min), suspended in RPMI containing 20% fetal bovine serum (Thermo Fisher Scientific), filtered on a 70 µm cell strainer and then used for subsequent analysis or frozen in PSC Cryomedium (Life Technologies).

**Figure 2.**
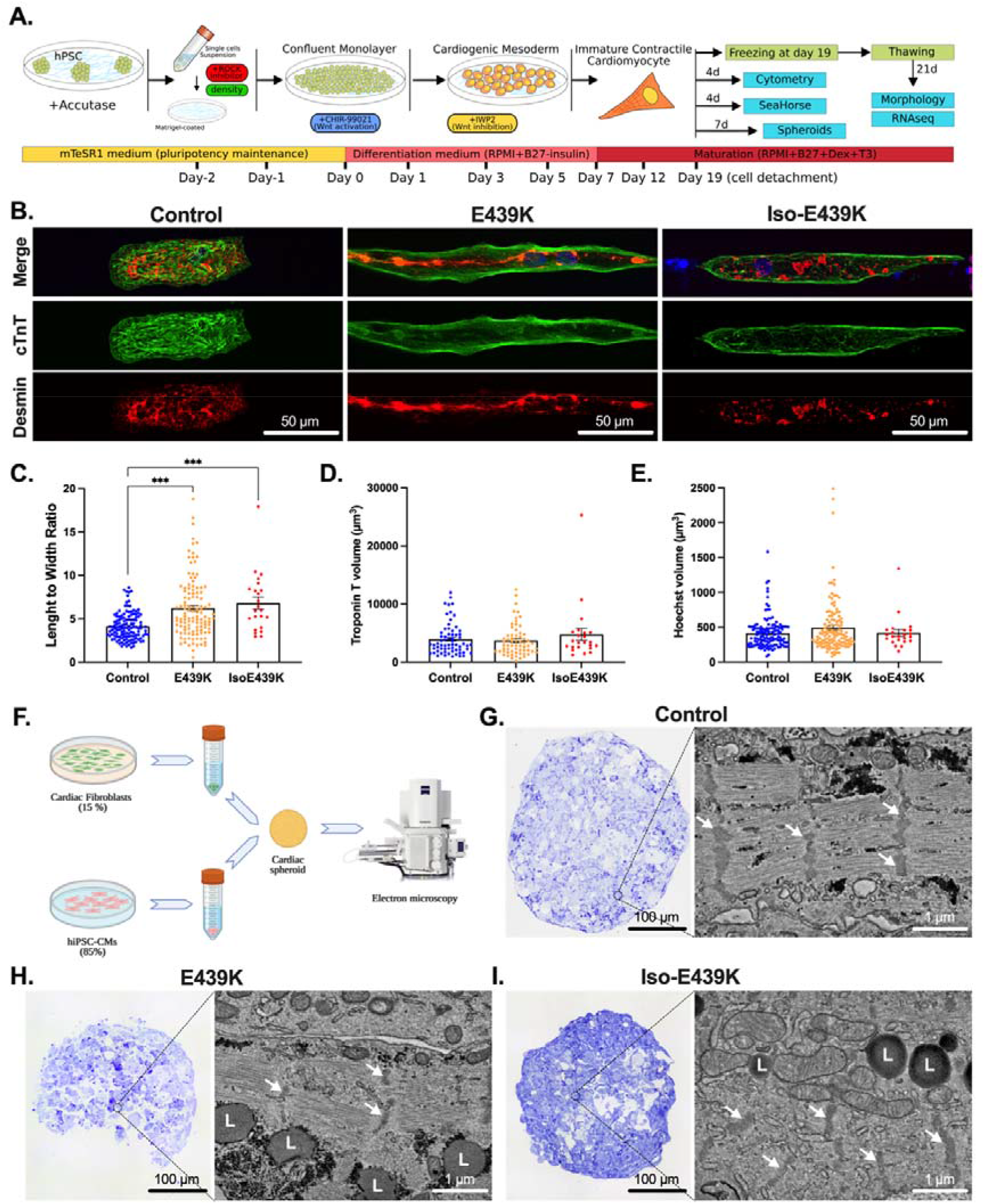
Morphological characterization of iPSC-derived cardiomyocytes carrying the *DES*^E439K^ mutation. (A) Schematic overview of the cardiac differentiation protocol from iPSC. After detachment or freezing at day 19, the cardiomyocytes are cultured for 4, 7 or 21 more days and analyzed. (B) Cardiac troponin T (Green) and Desmin (Red) immunostaining of cardiomyocytes cultured on gelatin micro-patterns (20 µm-wide lines) for 21 days after thawing. Nuclei were counterstained with Hoechst (blue). Note the marked disorganization of the sarcomeres and the accumulation of desmin in the cytoplasm of mutant cardiomyocytes. (C-E) Cardiomyocyte length-to-width ratio (C), cardiomyocyte volume (D, troponin T staining) and nuclei volume (E, Hoechst staining) were measured on cardiomyocytes after 21 days of culture on gelatin micro- patterns using ImageJ software. Values are given as means ± SEM. ***, p < 0.001. (F) Schematic representation of the production of a cardiac spheroid model used for electron microscopy analysis. (G-I) Bright-Field microscopy images of Toluidine blue-stained semithin sections and representative field emission scanning electron microscopy (FE-SEM) images from the full mapping of ultrathin sections of cardiac spheroids. White arrows indicate Z-lines; L, lipid droplets.

### Generation of cardiac spheroids

3D cardiac spheroid was generated using primary human cardiac fibroblasts (NHCFV, Lonza) cultured in FGM3 (Lonza) and iPSC-CM detached 19 days after the initiation of differentiation. Appropriate amounts of fibroblasts (1500/Spheroid) and iPSC-CM (8500/Spheroid) are mixed and seeded in 96-well spindle-shaped plate (S-BIO) in RPMI (Thermo Fisher Scientific) containing 20% of fetal bovine serum (Thermo Fisher Scientific). Medium was changed the day after, and every two days thereafter, with maturation medium. Spontaneously contractile spheroids were manually counted at day 7.

### Immunostaining and morphological analysis

Frozen iPSC-CM were thawed, centrifuged at 200xg to remove freezing medium and directly seeded on the coverslips micropatterned with gelatin lines (see Supplemental methods). They were then cultured 20 days in maturation medium, washed with PBS, fixed with 4% PFA for 5 min and permeabilized in cold methanol (-20°C) or 0.1% Triton prior to immunostaining. Paraffin-embedded human tissue sections were deparaffinized in successive bathes of xylene and ethanol. Antigenic epitopes were unmasked in citrate buffer pH 6.0 at 95°C for 30 min. Sections were incubated with 1×TrueBlack Lipofuscin Autofluorescence Quencher (Biotium) in 70% ethanol for 30 s at room temperature. Primary antibodies were incubated overnight at 4°C and secondary for 1 hour at room temperature diluted in PBS containing 2% of bovine serum albumin (see Supplemental Table 1 and 2). Slides were mounted with mounting medium containing Mowiol (Merck). Experiments were performed 3 times for Control-CMs (Control 1 and 2), patient E439K- CMs (clones c2 and c15) and isoE439K-CMs. At least 15 desmin-positive cardiomyocytes were imaged from each experiment with an inverted confocal microscope (Leica TCS SP5 AOBS, IBPS imaging platform). All images were acquired with a Z-step of 0.5 µm (image size 512x512, magnification 63x). The morphological analysis was performed using ImageJ and the 3D Manager plugin to measure length-to-width ratio and volumes of cells and nuclei.

### Western Blot

Human frozen cardiac tissue samples or iPSC-CMs pellets were solubilized in RIPA buffer (10 mM Tris HCl pH 7.4, 0.5% sodium deoxycholate, 1% NP40, 0.1% sodium dodecyl sulfate, 150 mM NaCl) in the presence of Protease Inhibitor Cocktail (Thermo Fisher Scientific 78425), Phosphatase Inhibitor Cocktail 2 (Sigma P5726) and Phosphatase Inhibitor Cocktail 3 (Sigma P0044). Each sample was incubated on ice for 30 min and centrifuged at 13,000g for 30 min at 4°C. Soluble proteins were then dosed using the Bicinchoninic Acid Kit (Merck) and diluted in Laemmli protein sample buffer. Proteins migrated into 4% to 20% mini-PROTEAN TGX stain-free gels (Bio-Rad) and transferred on polyvinylidene fluoride membrane (0.45 μm pore size) (Bio-Rad). After blocking in 0.1% PBS-Tween containing 5% non-fat milk, membranes were incubated overnight at 4°C with mouse monoclonal Total OXPHOS rodent WB cocktail (1:500, Abcam) or Rabbit monoclonal COXIV (1:1000, Cell Signaling). After 3 washes, membranes were then incubated with goat anti-rabbit IgG-HRP (1:20000, Sigma) or Goat anti-mouse IgG-HRP (1:10000, Sigma) for 1 hour at room temperature. Immunoreactivity was revealed by using Chemiluminescent Western Blot Reagents (Thermo Fisher Scientific) according to the manufacturer’s instruction. Images were acquired on a Bio-Rad Chemidoc system. Densitometry was normalized to the total protein loaded (stain free signal) for each lane using ImageJ.

### Sample preparation for ultrastructural analysis

Spheroids were fixed 1h with 1.5% glutaraldehyde, 1% paraformaldehyde and 0.04% Ruthenium Red in 0.1M sodium cacodylate buffer (pH 7.4), washed with the same buffer, incubated 1h with 0.2% Oolong Tea Extract (OTE) in 0.1M cacodylate buffer, and washed again. Samples were postfixed 1h with 1% osmium tetroxide, 1.5% potassium ferrocyanide in 0.1M cacodylate buffer, washed with deionized water, and embedded with 4% low melting point agarose. Samples were dehydrated with increasing concentrations of ethanol then acetone, infiltrated with resin (Agar 100 kit), and polymerized in standing gelatin capsules for 48 h at 60°C. Blocks were cut with an ultramicrotome (Ultracut UCT, Leica microsystems) according to planes of maximum diameter for the spheroids. 500nm semithin sections were deposited on glass slide, stained with Toluidine blue and observed with a Leica DM1000 microscope equiped with a XCAM 1080PHA (Touptek). Following 70-80nm ultrathin sections were deposited on 5 mm square silicon wafers or 150 mesh copper grids and were contrasted 15 min with 2.5% uranyl acetate. Wafers were stuck on aluminum stubs and observed with a Field Emission Scanning Electron Microscope (GeminiSEM 500, Zeiss), operating in high vacuum, at 1.5kV with the high current mode and a 30μm aperture diameter, and around a 2mm working distance. Two in-column detectors were used to separately collect secondary and backscattered electrons (filtering grid at 600V). To obtain transmission electron microscope-like images, those 2 channels were mixed and LookUp Table was inverted. Automated acquisitions were performed using Atlas 5 (Fibics), with a pixel dwell time of 6.4μs, a line average of 5, a pixel size of 8nm, an image definition of 5kx5k pixels, and on overlap of 15% between images forming a mosaic after stitching. Grids were observed at 80kV with a LaB_6_ JEM 2100 HC transmission electron microscope (Jeol) equiped with a side mounted Veleta CCD camera driven by iTEM software (Olympus). Images were recorded with an exposure time of 750ms and an image definition of a 2kx2k pixels.

### RNA sequencing

Frozen iPSC-CM from Control 1 and Clone c15 were thawed, centrifuged at 200xg to remove freezing medium and directly seeded on Matrigel-coated 12 well plates. They were then cultured 21 days in maturation medium, detached using TrypLE Select 1X, pelleted and flash frozen in liquid nitrogen. RNA was purified using Qiagen RNeasy microkits Plus (Qiagen). RNA was quantified using the Quant-iT RiboGreen RNA Assay Kit (Thermo Fisher Scientific) and quality control performed on a Bioanalyzer (Agilent), prior to mRNA library preparation using the Single Cell/Low Input RNA Library Prep Kit for Illumina® (New England Biolabs). Libraries were sequenced on an Illumina HiSeq 2500 V4 system. Sequencing quality control was performed using Sequence Analysis Viewer and FastQ files were generated on the Illumina BaseSpace Sequence Hub. Transcript reads were aligned to the hg18 human reference genome using Salmon v1.9.0^35^. Import and summarize transcript-level abundance to gene-level was performed with tximport package^36^. Quality control of the alignment was performed via MultiQC v1.4 . Finally, counts were normalized as counts per million. Differential gene expression analysis was performed with the DESeq2 R package^37^. Principal Component Analysis (PCA) was performed with factorMineR package^38^ and factoextra package^39^. The differentially expressed genes with adjusted *p* values false discovery rate (FDR)≤0.05 were subjected to GSEA with GO database using the ClusterProfiler R package^40^. The density plots were produced using the ggplot2 R package^41^. Weighted correlation network analysis (WGCNA) was applied on normalized data recovered from DESeq2 analysis. Pathways associated with each WGCNA modules were determined using METASCAPE web application^42^. WGCNA module genes were also submitted to protein-protein interactions using STRING web software (https://string-db.org/), and to MCODE analysis^43^ using Cytoscape software^44^. The heatmap was, performed with the R package pheatmap (version 1.0.12). All analysis was performed in R (v.4.2.0).

### PCR, 3D-PCR, cloning and sequencing

For these experiments, iPSC-CMs monolayer was dissociated using TrypLE Select 1X. A fragment of *MT-COI* (mitochondrial cytochrome c oxidase subunit I) gene was amplified by PCR and by differential DNA denaturation polymerase chain reaction (3D-PCR). This technique relies on the fact that heat denaturation of AT-rich DNA occurs at a lower temperature compared to GC-rich DNA^45,46^. Initial PCR conditions were a first cycle: 95 °C for 5 min, followed by 40 cycles (95 °C for 30 s, 60 °C for 30 s, and 72 °C for 2 min), and finally 10 min at 72 °C with the following primers, *MT-COI-*ext5-5′- GCGGTTGACTATTCTCTACAAACCACAAA-3’ and *MT-COI*-ext3- 5′GGGGGTTTTATATTGATAATTGTTGTGATGAAA-3’. 3D-PCR was performed with 1:50 of the first round PCR products. 3D-PCR was performed on an Eppendorf gradient Master S programmed to generate a 77-85°C gradient of denaturation temperature. The reaction parameters were a first cycle: 77 to 85 °C for 5 min, followed by 40 cycles: 77-85 °C for 30 s, 60 °C for 30 s, and 72 °C for 2 min, and finally 10 min at 72 °C. Primers used were *MT-COI-*int5-5’- CGTTATCGTCACAGCCCATGCATTTGTAA-3’ and *MT-COI-*int3-5′-GAGGAGACACCTGCTAGGTGTAAGGTGAA-3’. 3D-PCR products were purified from agarose gels (NucleoSpin Gel and PCR Clean-up, Macherey-Nagel) and ligated into the TOPO TA cloning vector (Invitrogen). About 100 colonies were sequenced.

### Real-time PCR

All RNA was extracted from the cells using the RNeasy Plus minikit (Qiagen) and cDNA synthesis was performed using QuantiTect reverse transcription kit (Qiagen) from ∼1 µg of RNA. Expression of APOBEC3 genes was assayed by real-time PCR based on TaqMan (Applied Biosystems) along with *RPL13A* as a reference gene. Primers used for the amplification of APOBEC3G were previously described^47^.

### Seahorse

20 days after the initiation of differentiation, iPSC-CM were dissociated using TrypLE Select 1X, seeded (25,000-50,000 cells/well) on Matrigel coated test plates (Extracellular Flux Assay Kit, Agilent) and cultured 4 days in maturation medium prior to measurement. The assay was performed in test medium composed of bicarbonate-free RPMI (pH=7.4) supplemented with glucose (4.5 g/L), sodium pyruvate (100 µM) and glutamine (200 µM). Cells were washed twice and pre-incubated at 37°C without CO_2_ in 500 µL of test medium for 1 h before measurement. For the Mito Stress test, 50µL of stock solution of each drug were injected sequentially during the experiment. Stock solutions of oligomycin (10 µM), FCCP (10 μM) and rotenone + antimycin A (1 μM) were prepared in DMSO, stored at -20°C and new aliquots were thawed extemporaneously for each experiment. For normalization, OCR values have been divided by the number of cardiomyocytes per well as counted after immunostaining for cardiac troponin T. Calibration to correct for the plate effect^48^ has been done by dividing all the OCR values by the lowest OCR value of the plate after addition of Rotenone and Antimycin. Basal and ATP-linked respirations, Maximal and Reserve capacities and OCR/ECAR were calculated as described in^49^. The experiment was performed 5 times with Control-CMs (Control 1 and Control 2), E439K-CMs (clones c2 and c15) and 2 times with Iso- E439K-CMs.

### Isolation and transfer of extracellular vesicles containing mitochondria

The extracellular vesicles were obtained from Control-CMs-conditioned medium based on differential centrifugations. Floating cells, cell debris and apoptotic bodies were removed by centrifugation at 1,000xg for 10 min. Pelleted vesicles were obtained by centrifugation of the supernatant at 10,000xg for 30 min at 4 °C. The quantity of extracellular vesicles was evaluated by measuring the protein concentrations that were measured using Bicinchoninic Acid Kit (Sigma). Cardiomyocytes or spheroids were treated with pelleted vesicles corresponding to 0.1 mg of proteins per 25 000 cells every two days from day 10 until day 24 after the initiation of differentiation.

### Statistical analysis

All experimental data are presented as mean ± standard error of the mean. Normality was verified using the test of Shapiro-Wilk and, when necessary, non-parametric tests were used. Statistical significance between two groups was determined using the Mann Whitney test. When multiple comparison was necessary, One-Way ANOVA combined with Dunnett’s multiple comparison tests or Kruskal-Wallis test combined with Dunn’s multiple comparison tests were used. A p-value of less than 0.05 was considered statistically significant. Data were analyzed and presented using GraphPad Prism 8. RNA-sequencing data that support the findings of this study have been deposited in Gene Expression Omnibus (GEO) repository with the accession codes XX.

## Results

### Morphological and mitochondrial alterations in iPSC-derived cardiomyocytes from the *DES*^E439K^ patient successfully recapitulate cellular defects of MFM1 hearts

To investigate the pathophysiological mechanisms of this heterozygous familial *DES*^E439K^ mutation, we generated two iPSC clones from the peripheral blood mononucleated cells (PBMC) of the patient CIII using a non-integrating episomal plasmid-based reprogramming strategy. Then, cardiomyocytes from *DES*^E439K^ iPSC (E439K-CMs) and two healthy control iPSC (Control-CMs) were generated using the biphasic activation/inhibition of the Wnt pathway differentiation protocol and detached or frozen at day 19 from first induction of cardiac differentiation (**Figure-2A**). Cardiomyocytes obtained from these iPSC clones were compared with cardiomyocytes from an isogenic pair carrying the same mutation which has been introduced by CRISPR/Cas9 (Iso-E439K-CMs) in one healthy iPSC clone (**Figure-S1**). In addition, when applicable, results from cardiomyocytes derived from iPSC (IPSC-CMs) were confirmed with heart biopsies of suddenly died Index case CII of the same family harboring *DES*^E439K^ mutation, and post-mortem heart samples from five control healthy patients. Since the diagnosis of MFM1 is most often made by the observation of structural perturbations, we first conducted a series of immunostaining on iPSC-CMs cultured on gelatin micro-patterns (20 µm-wide lines) (**Figure-2B**). After thawing, cardiomyocytes were cultured for 21 days in a medium containing T3 hormone and dexamethasone which are known to improve cardiac maturation of iPSC-CMs^50^. In Control-CMs, sarcomeres, as visualized by troponin T staining, display an almost well-organized structure and are surrounded by a filamentous desmin network throughout the cytoplasm. In contrast, E439K-CMs as well as Iso-E439K-CMs are characterized by a strong disorganization of sarcomeres as well as accumulation and patchy aggregation of desmin within the cytoplasm (**Figure-2B**). These observations faithfully reproduce the structural defects of cardiomyocytes as detected in human cardiac biopsies of the *DES*^E439K^ patient after immunostaining for troponin T and desmin (**Figure-S2A**). To examine closely the morphological alterations of E439K-CMs and Iso-E439K-CMs compared to Control-CMs, we measured cardiomyocyte length-to-width ratio, as well as nuclei and cardiomyocyte volumes. The length-to-width ratio is significantly lower in Control-CMs as compared to E439K-CMs and Iso-E439K-CMs (4.13±0.14, n=128 for Control-CMs *vs* 6.22 ± 0.32, n=118 for E439K-CMs or 6.81 ± 0.70, n=23 for Iso-E439K-CMs, *p*-value<0.001) (**Figure-2C**). Of note, high variability of length-to-width ratio was observed in mutant cardiomyocytes, which is consistent with the phenotype observed in DCM patients^51^. We validated the relevance of this parameter by showing that the length-to-width ratio of cardiomyocytes in biopsies was significantly higher in the heart of *DES*^E439K^ patient compared to healthy hearts (5.21±0.14 for Control *vs* 7.00±0.59 for *DES*^E439K^, *p*=0.003) **(Figure-S2B**). No differences were found in the volumes of cells and nuclei between healthy and mutated cardiomyocytes, as calculated on the basis of the Troponin T and Hoechst staining respectively (**Figure-2D-E**), indicating that the change of cell shape is not due to a hypertrophic mechanism.

We pursued the investigation of cellular morphological alterations in mutant cardiomyocytes by ultrastructural analysis using electron microscopy. To increase the physiological relevance of our results, cardiac spheroids, providing a tissue-like 3D model^52^, were generated using primary cardiac fibroblasts and iPSC-issued cardiomyocytes. Seven days after detachment (see schematic protocol in **Figure-2A**), fibroblasts and cardiomyocytes were mixed in a ratio 1.5:8.5 and cultured 7 more days in PrimeSurface 96M plate to favor their self-aggregation to spheroids (**Figure-2F**). For a global analysis of the cardiomyocyte ultrastructure under the 3 experimental conditions, a full mapping of each spheroid (sectioned at its maximum diameter) was carried out using field emission scanning electron microscopy (FE-SEM) and automated image acquisition. Electron microscopy studies disclosed a particularly well-organized ultrastructure of Control-CMs in terms of sarcomere organization and Z-line alignment (**Figure-2G**). In contrast, *DES*^E439K^ mutation resulted in a strong disorganization of sarcomeres which are less abundant and often misaligned between each other (**Figure-2H-I**), as revealed by shifted Z-lines (white arrows).

As shown in **Figure-2H-I**, we also noticed a strong cytoplasmic accumulation of lipid droplets in E439K-CMs and Iso-E439K-CMs as compared to Control-CMs, strongly suggesting that cellular metabolism may be affected in mutant cardiomyocytes. Following this observation, we investigated whether desmin mutated cardiomyocytes recapitulate mitochondrial abnormalities known to occur in hearts of MFM1 patients. For this, COXIV immunostaining was firstly assessed in E439K-CMs and Iso-E439K-CMs and compared to Control-CMs cultured on gelatin micro-patterns for 21 days after thawing. While COXIV-positive mitochondria were detected in Control-CMs, in particular at the extremities of cardiomyocytes and in the perinuclear area, this marker was not detectable in E439K- CMs and Iso-E439K-CMs (**Figure-3A**). Consistent with this observation, the expression of mitochondrial respiratory chain proteins, including respiratory complex I marker NDUFB8 an complex IV marker COXIV, was found decreased in mutated cardiomyocytes as compared to Control- CMs by Western blot analysis (**Figure-3B**). Once again, the mitochondrial defects detected in iPSC-derived mutated cardiomyocytes were confirmed in the heart of the desmin mutated patient since COXIV labeling (**Figure-S2A**) and Western blot analysis of NDUFB8 and COXIV markers (**Figure-S2C-E**) reveal similar decreased expression of these proteins in the diseased heart as compared to healthy ones. We then looked for an explanation for the decreased expression of respiratory mitochondrial proteins in desmin mutated cardiomyocytes. We first demonstrated that decreased expression was not due to a lower number of mitochondria in E439K-CMs. Microscopy and flow cytometry analysis of fluorescent MitoTracker-labeled mitochondria revealed no significant difference in their cytoplasmic distribution and quantity between mutated and control cardiomyocytes (**Figure-3C-D**). To determine whether this decrease was reflecting abnormal mitochondrial structure associated with the *DES*^E439K^ mutation, transmission electron microscopy (TEM) analysis was performed according to a classification of mitochondria into three types, *i.e.* normal, vesicular and swollen^53^ (**Figure-3E)**. Normal mitochondria are identified based on an appropriate architecture of cristae while vesicular mitochondria present enclosures of the internal membrane which creates several compartments within mitochondria. Finally, swollen mitochondria are identified by less dense staining of their matrix, expanded matrix space and fewer cristae. The quantification of mitochondria using this classification revealed a drastic increase of vesicular and swollen mitochondria in E439K-CMs compared to Control-CMs (**Figure-3F**), indicating important structural deficiencies in mitochondria of mutated cardiomyocytes that could explain the decrease in respiratory mitochondrial protein levels. Taken together, our results provide evidence that cardiomyocytes derived from iPSC of the *DES*^E439K^ patient successfully recapitulate crucial structural changes and mitochondrial alterations that are observed in biopsies of patients with the MFM1-related DCM, supporting the relevance of this cell model to study the pathology.

**Figure 3.**
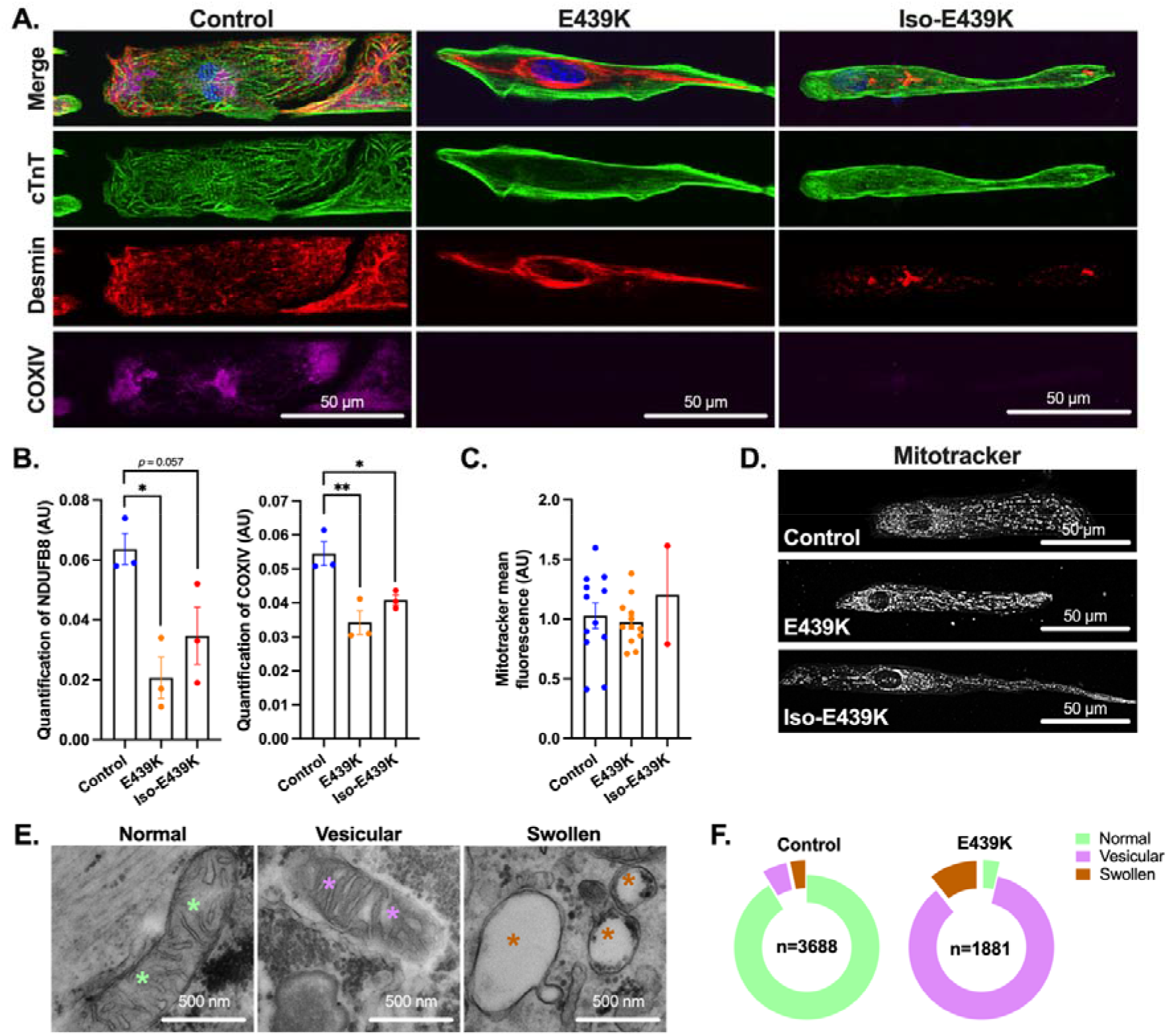
Structural and mitochondrial abnormalities in cardiomyocytes with the *DES*^E439K^ mutation. (A) Cardiac troponin T (Green), Desmin (Red) and COXIV (purple) immunostaining of cardiomyocytes cultured on gelatin micro-patterns (20 µm-width lines) for 21 days after thawing (see Figure 2A). Nuclei were counterstained with Hoechst (blue). Note the absence of COXIV labelling in E439K-CMs and Iso-E439K-CMs. (B) Quantitative analysis of Western Blot of NDUFB8 and COXIV showing the marked decrease of mitochondrial proteins in E439K-CMs and Iso-E439K-CMs compared to Control-CMs. Results are presented after normalizing to Stain-Free total protein profile. Results are expressed as mean values ± SEM. *, p < 0.05, **, p < 0.01. (C) Quantification of mitochondria in cardiomyocytes by cytometry using MitoTracker, indicating close levels of mitochondria between mutant and control cardiomyocytes. Values are given as means ± SEM. (D) Visualization of mitochondria using the mitochondria-specific fluorescent probe MitoTracker in iPSC-derived cardiomyocytes, indicating no significant differences in the cytoplasmic distribution of mitochondria in mutant cardiomyocytes. (E) Representative transmission electron microscopy (TEM) images of three types of mitochondria, *i.e.* normal (green asterisks), vesicular (purple asterisks) and swollen (brown asterisks), observed in iPSC-derived cardiomyocytes. (F) Ring chart showing the proportions of the 3 different types of mitochondria in E439K-CMs (n=1881: 3.6% normal, 85.8% vesicular and 10.6% swollen mitochondria) and Control-CMs (n=3688: 91% normal, 5.4% vesicular and 3.6% swollen mitochondria) revealing a switch of mitochondria from the normal to the vesicular type in E439K-CMs.

### iPSC-derived cardiomyocytes from the *DES*^E439K^ patient suffer severe mitochondrial dysfunction

To investigate whether metabolism is altered in *DES*^E439K^ cardiomyocytes derived from iPSC, we first performed RNA sequencing to compare the transcriptional expression profiles of iPSC-derived E439K-CMs and Control-CMs cultured during 21 days after thawing. According to the principal component analysis (PCA), the samples of both groups are well segregated after the dimensionality reduction of gene expression (**Figure-4A**), indicating that desmin mutation induces strong changes in the transcriptome of the cardiomyocytes. To obtain a deeper insight into the impacted biological processes in cardiomyocytes carrying the *DES*^E439K^ mutation, a gene set enrichment analysis (GSEA) was performed to determine the sets of genes differently expressed between E439K-CMs and Control- CMs (**Figure-4B**). This analysis reveals that E439K-CMs are highly enriched in sets of up-regulated genes related to cell division, DNA repair and acute inflammation, consistent with an adaptive response of cardiac cells to DNA damage, inflammation and oxidative stress. In addition, E439K-CMs were found to exhibit less enrichment for a set of down-regulated genes related to the sarcomere organization, heart contraction, mitochondrial electron transport and respiratory chain complex I confirming the impact of the *DES*^E439K^ mutation on the sarcomeric structure and mitochondrial functionality in cardiomyocytes. To further analyze the RNAseq data, we performed a non-supervised Weighted Gene Correlation Network Analysis (WGCNA) to highlight clusters of highly correlated genes. Two clusters (Blue and Red) were determined as highly statistically significant (**Figures-4D** and **S3A** for blue, and **Figure-S3B-D** for red cluster). We decided to focus on the blue cluster because the enrichment analysis of the genes that are co-expressed in this cluster revealed several sets of genes related to mitochondria (*i.e.* oxidative phosphorylation, mitochondrial organization, mitochondrial gene expression, respiratory electron transport, proton transmembrane transport). The expression of the majority of genes within these sets (**Figure-4E-G**) are strongly downregulated in E439K-CMs, supporting that mitochondrial disruption is an important hallmark of the patient harboring the *DES*^E439K^ mutation.

**Figure 4.**
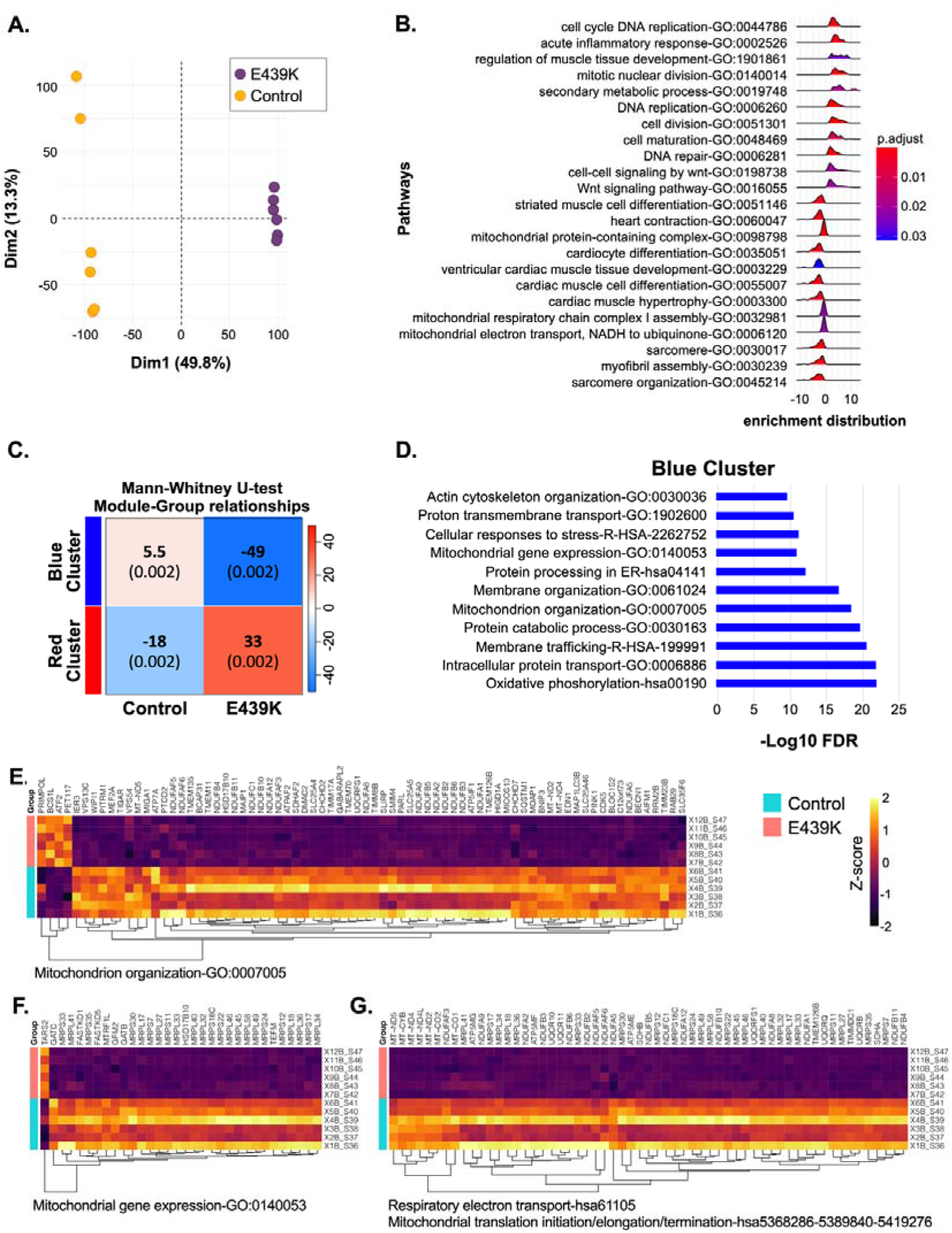
Transcriptomic analysis of E439K-CMs compared to Control-CMs reveals a strong signature of mitochondrial impairment. **(A)** Principal component analysis (PCA) of gene expression of the different samples from E439K-CMs (n=6) and Control-CMs (n=6). **(B)** Gene Set Enrichment Analysis (GSEA) of E439K-CMs compared to Control-CMs. **(C-G)** Non-supervised Weighted Gene Correlation Network Analysis (WGCNA) have detected two significant clusters of genes **(C)**. The molecular complex determination and enrichment analysis of the Blue cluster **(D)** reveals different sets of genes related to mitochondria **(E-G)**.

We then investigated the impact of mitochondrial gene and protein alterations on the respiratory activity measured with the Seahorse metabolic analyzer in desmin mutated cardiomyocytes 4 days after thawing (**Figure-5A-F**). Using the mito stress test, we found strong, statistically significant differences between Control-CMs and cardiomyocytes carrying the *DES*^E439K^ mutation (E439K-CMs and Iso-E439K-CMs) in all examined parameters. The mito stress test allows to assess in real-time the oxygen consumption rate (OCR) at the initial level as well as after the addition of inhibitors that target specific respiratory complexes (**Figure-5A**) to evaluate different parameters such as basal respiration, ATP-linked respiration, maximal respiration and reserve capacity (**Figure-5B-F**). Basal respiration is calculated from the OCR difference between the initial state and after complete blockage of the respiratory chain by addition of rotenone and antimycin A. As shown in **Figure 5B**, a decrease of basal respiration was detected in *DES*^E439K^ cardiomyocytes compared to Control-CMs. Addition of oligomycin, an ATP synthase inhibitor, allows to measure ATP-linked respiration, the consumption of oxygen specifically used for the biosynthesis of ATP. This parameter is lower in *DES*^E439K^ cardiomyocytes (**Figure-5C**), suggesting a lower involvement of mitochondria in ATP generation in the presence of the mutation.

**Figure 5.**
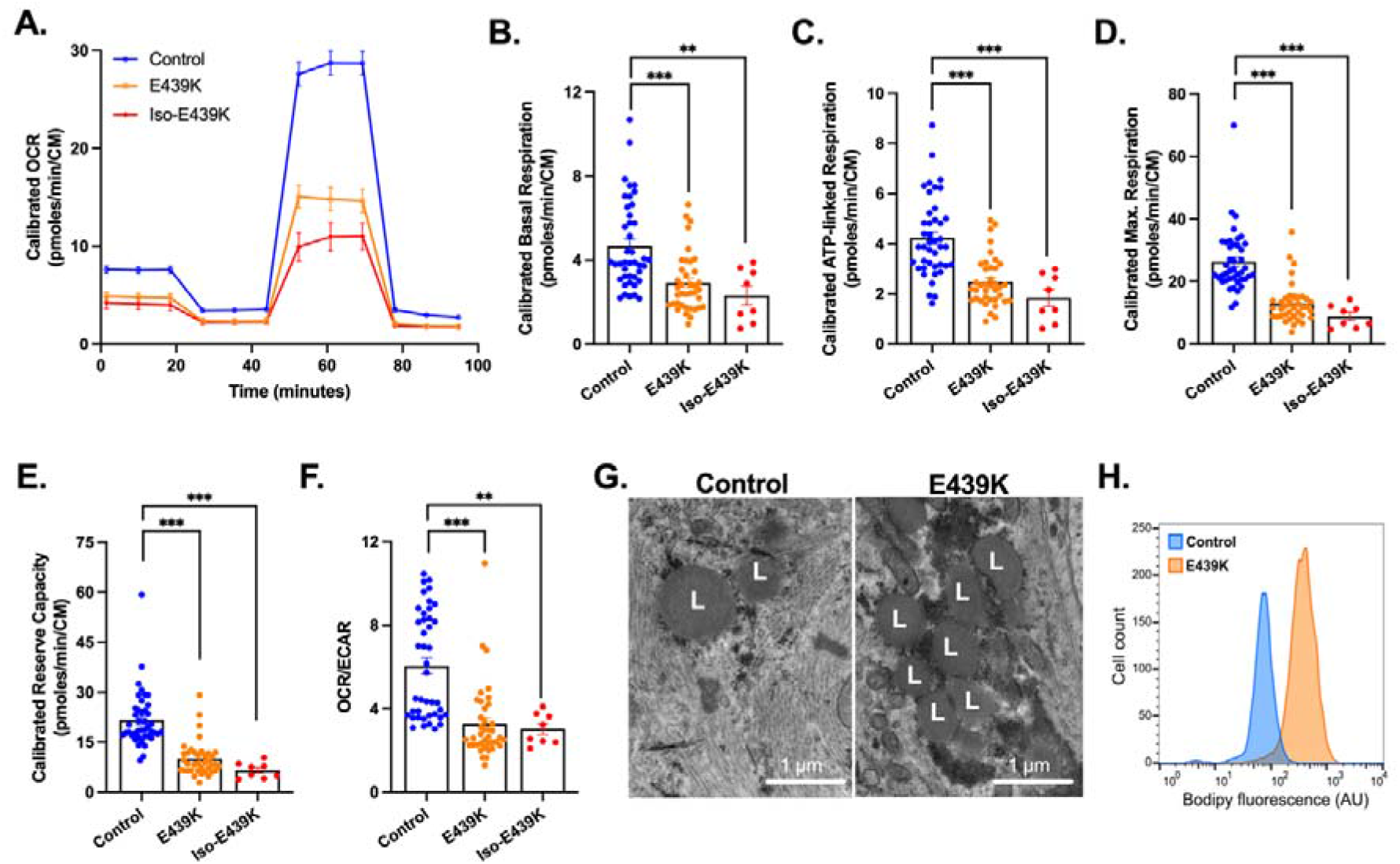
Decreased mitochondrial respiration and altered metabolic activity in cardiomyocytes with *DES*^E439K^ mutation. **(A)** Representative mitochondrial oxygen consumption rate (OCR) profiles in Control- CMs, E439K-CMs and Iso-E439K-CMs. OCR profiles are expressed as pmol O_2_/min normalized to the number of cardiomyocytes and calibrated to the lowest value of the plate after addition of rotenone and antimycin A. **(B)** Quantification of basal respiration, **(C)** ATP-linked respiration, **(D)** maximal respiration and **(E)** reserve capacity of cardiomyocytes. **(F)** Ratio of OCR to ECAR in iPSC-derived cardiomyocytes. Values are expressed as mean±SEM. **, *p*<0.01, ***, *p*<0.001. (G) Representative transmission electron microscopy images of lipid droplets (L) in iPSC-derived cardiomyocytes. (H) Flow cytometry analysis of cardiomyocytes after BODIPY staining demonstrating higher lipid levels in E439K-CMs.

Carbonyl cyanide-4 (trifluoromethoxy) phenylhydrazone (FCCP) is an uncoupling agent that collapses the proton gradient and disrupts the mitochondrial membrane potential. Addition of FCCP results in uninhibited electron flow through the electron transport chain, allowing to measure the maximal oxygen consumption rate. As shown in **Figure-5D**, the maximal respiration is lower in *DES*^E439K^ cardiomyocytes, indicating a lower total mitochondrial activity. The reserve capacity, defined as the OCR difference between basal respiration and maximal respiration, is also lower in *DES*^E439K^ cardiomyocytes (**Figure-5E**). Finally, to evaluate the role of glycolysis in producing energy/metabolites in cardiomyocytes, initial OCR was normalized to the initial extracellular acidification rate (ECAR). The ratio of OCR to ECAR, which indicates cellular preference for oxidative phosphorylation versus glycolysis^49^, was calculated for the different groups. As shown on **Figure-5F**, this ratio is lower in *DES*^E439K^ cardiomyocytes. Taken together, compared to Control-CMs, both types of mutant E439K-CMs have decreased mitochondrial respiration compensated by a higher glycolytic activity. In addition, electron microscopy analysis also revealed a strong cytoplasmic accumulation of lipid droplets in E439K-CMs and Iso-E439K-CMs as compared to Control-CMs (L, **Figure-5G**), that was confirmed by cytometry quantification of intracellular lipid using BODIPY, a fluorescent hydrophobic dye for lipids (**Figure-5H**). Taken in concert, these findings indicate that the metabolic activity of cardiomyocytes is altered when desmin is mutated.

### APOBEC3G is up-regulated and involved in the editing of released mitochondrial DNA (mtDNA) in iPSC-derived cardiomyocytes from the *DES*^E439K^ patient

To further improve our understanding of the mitochondrial perturbation in cardiomyocytes with the *DES*^E439K^ mutation, the mitochondrial DNA (mtDNA) was analyzed. It is known that different stimuli and/or cellular stresses may induce the release into the cytoplasm of mtDNA which in turn triggers DNA sensor molecules to induce an inflammatory response^54–56^, contributing to disease progression. We hypothesized a correlation between cell stress due to mitochondrial dysfunction and/or alteration of the cardiomyocyte structure, and the release of the mtDNA into the cytoplasm, where it gets edited. Using a differential DNA denaturation polymerase chain reaction (3D-PCR) we analyzed mtDNA editing on a fragment of mitochondria-specific *MT-COI* (mitochondrial cytochrome c oxidase subunit I) gene, in control (Control-CMs, n=5) and mutant cardiomyocytes (E439K-CMs, n=6 and Iso-E439K- CMs, n=2) (**Figure-6A**). A plasmid containing *MT-COI* DNA fragment and two cell lines (HeLa and 293T) were used as experimental controls (**Figure-6B**). 3D-PCR amplifies preferentially AT-rich DNA fragments that denature at a lower temperature than GC-rich DNA^45,46^. 3D-PCR was performed with a 77-85°C gradient in the denaturation temperature (Td). Interestingly, 3D-PCR products were recovered at a Td as low as 79.7°C for E439K-CMs and Iso-E439K-CMs while *MT-COI* plasmid set the threshold Td at 85°C (**Figure-6A-B**). We also noted that *MT-COI* DNA isolated from HeLa and 293T cells lines were detected with a Td of 82.7°C, which far exceeded the Td of the plasmid containing *MT-COI* DNA fragment (**Figure-6B**). These cell lines are perpetually in balance between fission and fusion of mitochondria which results in the release of mtDNA in the cytoplasm followed by its editing^57^. To explore editing at the molecular level, 3D-PCR products recovered at 79.7°C (E439K-CMs and Iso-E439K-CMs) and 80.7°C (Control-CMs) were cloned and sequenced. Extensive and monotonous G>A hyperediting was observed in *MT-COI* DNA from all three cardiomyocyte types (**Figure-6C**). Interestingly, the mutation frequency observed in E439K-CMs and in Iso-E439K- CMs (40-44 mutations in 248 bp; ∼17%) was higher than Control-CMs (35-39 mutations in 248 bp; ∼14%), demonstrating that the high mutation burden detected in mutant cardiomyocytes could be used as a marker for the detection of the mitochondrial network damage (**Figure-6D**).

**Figure 6.**
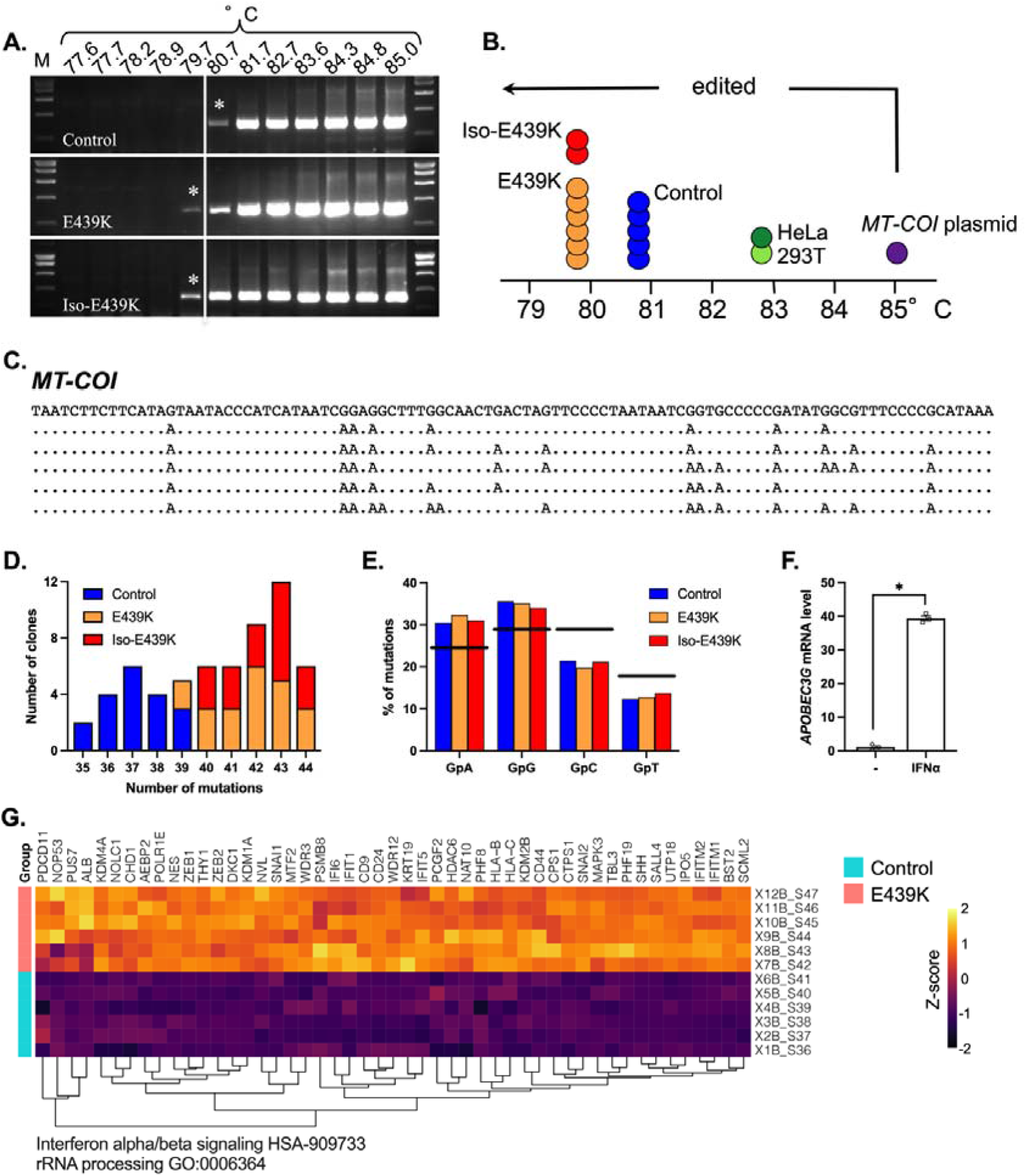
Editing of mtDNA in cardiomyocytes with the *DES*^E439K^ mutation using the *MT-COI* gene fragment. **(A)** mtDNA editing was analyzed using a denaturation temperature (Td) gradient across the heating block on first round PCR products. 3D-PCR recovered edited *MT-COI* DNA down to 79.7°C for E439K-CMs and Iso-E439K-CMs. The vertical white line indicates the threshold between Control-CMs, E439K-CMs and Iso-E439K-CMs 3D-PCR products in terms of the Td. M: molecular weight markers. Asterisks refer to the samples cloned and sequenced. **(B)** Schematic representation of the Td for the last positive 3D-PCR amplifications for 6 E439K-CMs samples (orange circles), 2 Iso-E439K-CMs samples (red circles), 5 Control- CMs samples (blue circles), non-transfected HeLa (dark green circle) and 293T (light green circle) cells and cells transfected with a plasmid containing *MT-CO*I DNA fragment (purple circle). The arrow indicates the threshold Td (85°C) at which the samples are hypermutated. **(C)** A selection of hypermutated G>A edited E439K-CMs and Iso-E439K-CMs samples (Td=79.7°C). The sequences are given with respect to the plus or coding strands. Only differences are shown. All sequences were unique, indicating that they corresponded to distinct molecular events. **(D)** Frequency analysis of edited *MT-COI* fragment gene as a function of the number of edits per sequence at 79,7°C and 80.7°C for the E439K-CMs and Iso-E439K-CMs-derived and for the control-CMs derived 3D-PCR products respectively. The size of the columns indicates the combined numbers of sequences analyzed across the three samples. **(E)** Bulk dinucleotide context of *MT-COI* gene fragment and compared to the expected values. The horizontal line represents the expected frequencies assuming that G->A transitions were independent of the dinucleotide context and correspond to the weighted mean dinucleotide composition of the reference sequence. The blue (Control-CMs), orange (E439K-CMs) and red (Iso-E439K-CMs) bars represent the percentage of G>A transitions occurring within 5’GpN dinucleotides for the hypermutated sequences. Preferential mutations in 5’GpA and 5’GpG contexts correspond to an APOBEC3G signature. (F) Gene expression quantification of APOBEC3G in Control-CMs evaluated by RT-qPCR after treatment with IFNα (1000U/ml). **(G)** Molecular complex detection of weighted gene correlation network analysis (WGCNA) from RNAseq data of E439K-CMs and Control-CMs have detected sets of genes related to IFN IFNa/b and rRNA processing.

The 5′ dinucleotide context associated with editing was strongly in favor of 5′GpA and 5′GpG (**Figure-6E**), which is typical for polynucleotide cytidine A3 deaminases (APOBEC3), responsible for G>A (or C>U in the opposite DNA strand) editing. A3 enzymes (isoforms APOBEC3A to APOBEC3H) leave a telltale editing signature in DNA, namely they preferentially edit a cytidine residue in the context of 5′GpA (5’TpC in the opposite DNA strand) with the exception of APOBEC3G, which prefers 5′GpG dinucleotides (5’CpC in the opposite DNA strand)^46,58–60^. Therefore, at least APOBEC3G appears to be involved in mtDNA editing in iPSC-derived cardiomyocytes. Interestingly, our RNAseq data also showed an upregulation of the transcriptional expression of APOBEC3G in E439K-CMs compared to Control-CMs (Fold change is 3.95, adjusted *p* value=1.31x10^-13^).

Since some of these enzymes (APOBEC3A, APOBEC3G) could be up-regulated during inflammation^61^ we then verified by a supervised analysis of our RNAseq data, whether sets of genes related to inflammation are up or downregulated between Control-CMs and E439K-CMs. Interestingly, the red cluster of genes differently expressed in E439K-CMs (**Figure-6C)** is in particular characterized by the enrichment of the Interferon alpha/beta (IFNα/β) signaling set of genes (**Figure-6G**), suggesting that inflammation may be triggered in mutant cardiomyocytes. These data indirectly support that dysfunctional mitochondria of desmin-mutated cardiomyocytes could exacerbate deleterious inflammation processes by releasing their mtDNA in the cytosol. Finally, to confirm the RNA sequencing-based transcriptome analysis concerning the relation between inflammation signaling pathways and APOBEC3G, Control-CMs were incubated during 24 hours with IFNα. As a result, APOBEC3G transcripts were increased more than 34-fold in IFNα-treated iPSC-derived cardiomyocytes (**Figure-6F**). These data demonstrated that upregulation of APOBEC3G cytidine deaminase is possible in iPSC-derived cardiomyocytes under chronic inflammatory conditions. In addition, given that APOBEC3 enzymes are strictly cytoplasmic and do not localize within mitochondria^56^, *MT-COI* DNA editing process could occur exclusively in the cytoplasm. Taken together, these results strongly suggest that the release of mtDNA into the cytoplasm possibly occurs, is more important in *DES*^E439K^ cardiomyocytes (E439K-CMs and Iso-E439K-CMs) compared to Control-CMs, and may induce stress responses that aggravate cellular defects.

### Transfer of exogenous functional mitochondria restores the metabolism and contractile function in desmin-mutated cardiomyocytes

Given that mutant cardiomyocytes suffer from severe mitochondrial defects, we were prompted to test whether the transfer of exogenous healthy mitochondria could restore proper metabolism and contractile function in cardiomyocytes carrying the *DES*^E439K^ mutation. We expected that isolated mitochondria enter into targeted cells, integrate the recipient’s endogenous mitochondrial networks and restore functional defects in iPSC-derived mutant cardiomyocytes.

As a first attempt, mitochondria were isolated from Control-CMs and immediately transferred to E439K-CMs or Iso-E439K-CMs. To explore the efficacy of transfer, exogenous mitochondria were labeled with MitoTracker before isolation, and transferred mitochondria were measured by flow cytometry 24h after their addition to cardiomyocytes. These experiments confirmed that mitochondria were efficiently transferred to E439K-CMs as well as Iso-439K-CMs as shown by the detected MitoTracker-positive cell population (**Figure-S4A**). We then investigated the consequences of this treatment in the metabolic activity of desmin-mutated cardiomyocytes. We found that uptake of exogenous mitochondria is accompanied by a general increase of the OCR profiles indicating the re-establishment of mitochondrial respiration in mutant cells (**Figures-S4B-C**). This phenomenon did not occur when the cells were treated with mitochondria previously inactivated with paraformaldehyde (PFA), supporting the metabolic role of exogenous mitochondria in recipient cells. Nevertheless, we found that 24h of treatment did not result in a significant improvement of mitochondrial respiration parameters (**Figures-S4B-C**), indicating that a short-term treatment may not be sufficient to restore cardiomyocyte bioenergetics. We therefore decided to evaluate the long-term effects of this treatment on mitochondrial respiration capacity as well as contractility of mutant cardiomyocytes (**Figure-7A**). In view of the high number of cells required for the isolation of mitochondria, we decided to simplify the protocol by switching from isolated mitochondria to the preparation of mitochondria-containing extracellular vesicles (EV) from the conditioned cell culture media, based on a recent publication by Ikeda et al.^62^. Flow cytometry analysis confirmed that mitochondria-containing EVs pre-treated with MitoTracker can effectively mediate the transfer of mitochondria into iPSC-CMs (**Figure-7D**). Cardiomyocytes carrying the *DES*^E439K^ mutation were treated with EVs every two days beginning day 10 up to day 20 of cardiac differentiation, and then their function in terms of respiration and contractility was evaluated. The beginning time of treatment (10 days of differentiation, see schematic protocol in **Figure-2A**) was chosen to test the hypothesis that the treatment may counteract the progressive accumulation of abnormalities in mutant cardiomyocytes. Indeed, mutant cardiomyocytes do not display any defects at the early time of differentiation neither in the expression profile of the different surrogate cardiomyocyte markers nor in mitochondrial respiration capacity (**Figure-S5A-C**). As shown in **Figure-7B-C**, this chronic treatment over 10 days results in an increase of OCR profiles of E439K-CMs and Iso-E439K-CMs compared to non-treated mutant cardiomyocytes. More interestingly, basal respiration, ATP-linked respiration and maximal respiratory capacity were all improved in both mutated iPSC-CMs compared to non-treated mutant cardiomyocytes. This functional improvement is also correlated to an increase in the percentage of spontaneously contractile spheroids (**Figure-7E**) monitored using MUSCLEMOTION ^63^. Essentially, spheroids derived from mutant cardiomyocytes did not demonstrate any spontaneous beating if not treated with mitochondria. Taken in concert, these experiments, reveal that supply of mutant cardiomyocytes with functional mitochondria is sufficient to correct the functional defects of these cells and support the hypothesis that mitochondria abnormalities are the main responsible factor for the cellular defects behind the pathological symptoms found in the heart of desmin-mutated patients.

**Figure 7.**
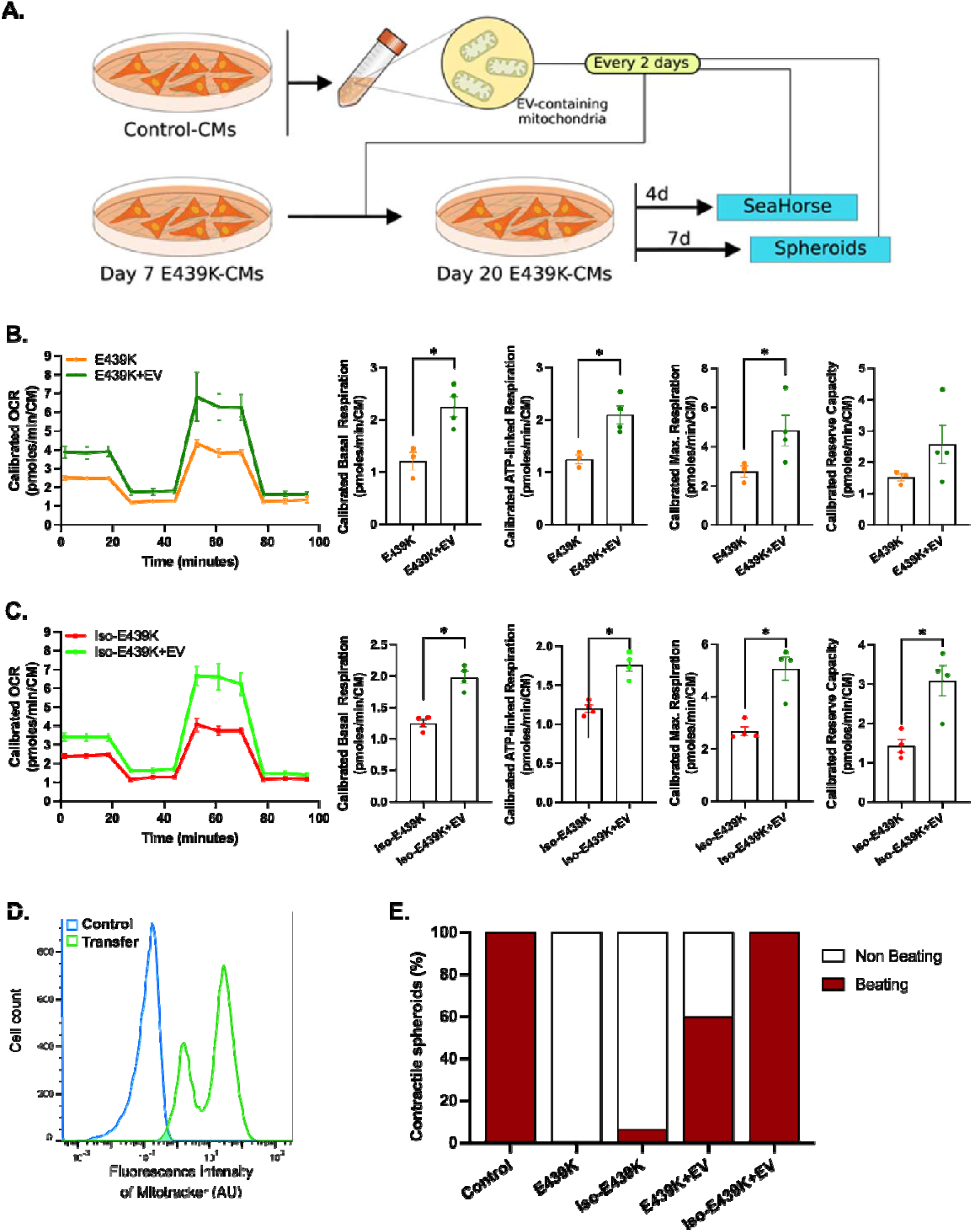
Long-term effects of the transfer of mitochondria on the function and respiration of cardiomyocytes with the *DES*^E439K^ mutation. (A) Experimental set-up. (**B-C**) Mitochondrial oxygen consumption rate (OCR) profiles and respiration parameters of the mito stress test of E439K-CMs (C) or Iso- E439K-CMs (D) untreated of treated with EVs-containing mitochondria (+EV). OCR profiles are expressed as pmol O_2_/min normalized to the number of cardiomyocytes and calibrated to the lowest value of the plate after addition of R/A. **(D)** Flow cytometry analysis of Control-CMs untreated (Control) or treated (Transfer) with extracellular vesicles (EV) from Control-CMs previously incubated with MitoTracker. **(E)** Percentage of contractile spheroids assessed by video microscopy

## Discussion

Emergence of cardiac dysfunction in desmin-related myofibrillar myopathies has been associated with sarcomeric disorganization and mitochondrial abnormalities in cardiomyocytes. However, the pathophysiological mechanisms triggered by desmin mutations are far from being understood. Data are coming mostly either from transgenic animals, which in many pathological cases respond differently as compared to humans, or from transiently transfected cells carrying human *DES* mutations which provide useful information, but often artificial due to the expression levels of transfected DNA. Otherwise, human biopsies from patients provide valuable information but are difficult to obtain and allow only descriptive evidence. We believe that further progress in this field necessitates relevant human cellular or tissue models that faithfully reproduce cardiac disease.

Recent advances in iPSCs seem to be a very promising response to overcome the limitations of previously studied models. Thus, we generated cardiomyocytes using iPSCs from a patient carrying the heterozygous *DES*^E439K^ mutation and from healthy donors as a control. To complete our analysis, we also created an isogenic pair of these control iPSCs in which the same mutation has been introduced. In an effort to increase physiological relevance, cardiomyocytes were either cultured on an anisotropic micropatterned surface (2D cultures), to obtain elongated and aligned cardiomyocytes, or as a cardiac spheroid (3D cultures), to create a micro-tissue. Using these cardiomyocyte preparations, we were able to show that (i) cardiomyocytes derived from iPSC of a *DES*^E439K^ patient successfully recapitulate the crucial structural changes and mitochondrial alterations observed in heart biopsies from this MFM1 patient, highlighting the relevance of these cell models to study the pathology; (ii) the observed severe structural and functional mitochondrial abnormalities represent a primary mechanism in the emergence of desmin-related cardiomyopathy in *DES*^E439K^ patients, since transfer of exogenous healthy mitochondria restores the metabolism and contractile function in mutant cardiomyocytes. Based on these findings, we propose that in cardiomyocytes, the primary disruption of the desmin network due to desmin mutations induces mitochondrial defects which, in turn, contribute to a vicious circle leading to cardiomyocyte dysfunction and subsequently to DCM.

To the best of our knowledge, our study is the first that combines efficient protocols to generate cardiomyocytes from hiPSC and maturation techniques to study the pathological phenotypes and mechanisms of desmin-related MFM. To date only one publication has reported the characterization of cardiomyocytes derived from iPSC from patients with desmin mutations. In this work, Tse HF *et al.*^64^ used for the first time iPSC-CMs derived from a *DES*^A285V^ patient affected by DCM. They showed that mutant iPSC-CMs displayed structural and functional abnormalities, namely a disorganized ultrastructure (alteration of intercalated discs, streaming of Z-disks, absence of I bands, accumulation of granulofilamentous material reminiscent of desmin aggregation) and an altered response to isoproterenol (lower rate of calcium re-uptake and a slower spontaneous beating rate). In this study the cardiomyocytes were obtained by a protocol based on the co-culture of embryoid bodies with endoderm-like cells, dissected and dissociated 6 weeks after the initiation of differentiation and analyzed as single cells or cell clusters. This approach clearly limited the quantity and purity of obtained cardiomyocytes. These limitations did not affect the study since, according to authors, the model was mainly used to assess genetic basis of clinically observed DCM by providing histological and functional confirmation. However, this model would not be optimal for further analysis of pathological cell mechanisms. In parallel, Brodelh and collaborators published a series of studies where healthy hiPSC-CMs were transiently transfected with different mutants of desmin^20,65–68^. Unfortunately, the mitochondrial function and/or phenotype was not characterized in these cardiomyocytes, probably because the efficiency of transient transfection was not sufficient to allow such analyses.

In our approach, after differentiation based on the monolayer GiWi protocol^34^, hiPSC-CMs were 2D- seeded on a gelatin micropattern and treated with T3 and Dexamethasone, to improve maturation and analyze their morphology. Moreover, we generated a 3D spheroid model that is expected to better mimic the organization of the native cardiac tissue^52^. Our results were further validated through the generation of an isogenic pair by introducing the *DES*^E439K^ mutation into a control hiPSC clone, which rigorously establishes that this heterozygous desmin point mutation is at the origin of the observed pathogenic phenotype. Finally, by comparing with samples from a patient carrying the same mutation, we were able to show that our cellular models successfully mimic several features of the MFM cardiac phenotype, as discussed below. All these specificities of our study carry out an important step forward into the comprehension of the pathophysiological mechanisms that underlie desmin-related genetic-driven DCM.

This work highlights the structural and functional alteration of mitochondria as the hallmark and primary defect of *DES*^E439K^ cardiomyocytes. Even if the link between desmin and mitochondria is not perfectly understood, it is well established that one common trait in patients affected by MFM resides in defective mitochondria^69^. At least 8 different mutations of desmin (*i.e.*, *DES*^R16C^, *DES*^E114del^, *DES*^A213V^, *DES*^K239f*s*242^, *DES*^A357P^, *DES*^E245D^, *DES*^R350P^ and *DES*^K449T^) have been directly linked to mitochondrial dysfunctions in human tissue samples^16,18,19,70,71^. In these studies, detrimental effects of desmin mutations on the mitochondrial content and biogenesis, morphology and maximal rate of respiration, have been observed. Moreover, alterations in mitochondrial organization and architecture also described with other genes related with MFM such as α-B-crystallin^72^, myotilin^71^ and ZASP^71^. Our study has strengthened the idea that mitochondrial disruption plays an important role in pathophysiology of desmin-related MFM. Indeed, our results show that the *DES*^E439K^ mutation is detrimental for proper mitochondria structure, respiratory capacity as well as mtDNA integrity. As shown in **Figure-3C-D**, the number of mitochondria was found similar between control and mutant cardiomyocytes in culture, suggesting that their biogenesis is not affected by desmin mutation. However, a remarkably high percentage (>90%) of abnormal vesicular and swollen mitochondria was observed in the *DES*^E439K^ iPSC-derived cardiomyocytes compared to control cardiomyocytes which, in the contrary, displayed a normal mitochondrial morphology in their majority (>90%) (**Figure-3E-F**). This result is consistent with the idea that the desmin network playing an important role in the structural integrity of mitochondria^11^. Recently, a direct interaction between desmin and mitochondrial proteins including VDAC, Mic60 (a component of the MICOS complex) and ATP synthase was demonstrated by Dayal and colleagues^26^. The authors suggest that these interactions should participate in mito-protection at different levels. Desmin was also found tightly associated with plectin^21^ and appeared to regulate, at least indirectly, the function of VDAC^25^. In desmin knockout mice, Kay et al.^73^ described two populations of mitochondria with the emergence of often enlarged subsarcolemmal mitochondria disconnected from myofibrils. This is reminiscent of our observation regarding the change in proportions of different types of mitochondria between E439K-CMs and Control-CMs, suggesting that both loss and mutations of desmin lead to similar mitochondrial alterations. However, the question remains as to how the interaction between desmin and mitochondria regulates their structure and function, such as size or positioning, membrane potential, respiration and ATP production.

As we demonstrated in **Figure 3**, morphological alteration is accompanied by decreased expression of mitochondrial respiratory complex I marker NDUFB8 and complex IV marker COXIV, as evidenced by Western blot analysis. This phenotype is also described in skeletal muscles of hetero- and homozygous knock-in mice carrying the R349P desmin mutation^19^. Moreover, the presence of histochemically COX-negative (COX deficient) muscle fibers within muscle biopsies is considered to be a diagnostic hallmark of mitochondrial disease^74^. Interestingly, at variance with Western blots, immunolocalization experiments showed the presence of completely COX-negative fibers in mutant cardiomyocytes. One explanation for this discrepancy could be a possible defect in the assembly of the complex IV in mutant cardiomyocytes; another possibility would be the masking of antibody’s epitopes due to potential post-translational modification induced by pathological condition. RNA-seq data also confirmed in E439K-CMs the decreased expression of NDUFB8 and COXIV, as well as of many other genes related to mitochondria, *i.e.* involved in oxidative phosphorylation, mitochondrial organization, respiratory electron transport, proton transmembrane transport, and mitochondrial gene expression. Consistent with this observation, we demonstrated that *DES*^E439K^ induces a strong decrease of the mitochondrial respiratory function (see **Figure-5**). Basal and ATP-linked respirations, as well as maximal and reserve capacities are decreased in mutant cardiomyocytes. The overall decrease of these parameters, especially the maximal capacity, seems not to be related to a decrease in mitochondrial content, which was also observed previously in desmin knockout mice^11^. It was stated that a change in basal respiration coupled to a change in ATP-linked respiration and maximal capacities can be due to an adaptation of mitochondria to altered ATP demand of the cardiomyocyte^75^ rather than to an intrinsic dysfunction of mitochondria. Commonly, mitochondrial dysfunctions should be determined if the decrease of basal and ATP-linked respiration is not coupled to a change in maximal capacity^75^. Thus we cannot exclude that the observed changes in mitochondrial respiratory parameters may underlie changes in the global energetic characteristics of the cardiomyocyte^49^. A change in the metabolic status could also explain the observed changes in the OCR/ECAR ratio that can be related to an increase in anaerobic glycolysis *versus* oxidative phosphorylation^49^. In agreement with this hypothesis, we also observed increased expression of genes involved in lipid beta-oxidation and a strong increase in the presence of lipid droplets within the cytoplasm of cardiomyocytes (see **Figure-5G**). Indeed, neutral lipids can be accumulated in cardiomyocytes because of an alteration in β- oxidation^76^. A recent publication highlighted a reconfiguration of the mitochondria-related metabolic pathways including fatty acid transport, activation, and catabolism^77^ that is attributed, secondary, to defects in mitochondria. To test this hypothesis, the mito stress test could also be performed on isolated mitochondria instead on intact cells. Similar experiments have been performed on mitochondria isolated from the heart of control and desmin-null animals^11^ and did not reveal any differences in respiration rates, while a significant difference in V*max* of oxygen consumption was noted in desmin-null intact cardiomyocytes.

Another interesting result was that the majority of the sets of genes displaying an increased expression in E439K-CMs are related to cell division, DNA repair and acute inflammation, indicating that an adaptive response to DNA damage and inflammation is possibly induced by desmin mutation. We further investigated this possibility. It is known that, while most damaged mitochondria following a cellular stress are removed by autophagy, some mtDNA fragments clearly find their way to the cytoplasm where they act as a danger signal and can trigger cytoplasmic DNA sensor molecules to induce inflammatory responses^78^. In fact, it was recently reported that mtDNA induces Toll-like receptor 9-mediated inflammatory responses in cardiomyocytes leading to myocarditis^79^. Similar mechanisms might play a role in other chronic inflammatory diseases or after severe bodily injury. On the other hand, APOBEC3 enzymes hypermutate (edit) and initiate catabolism of cytoplasmic mtDNA, acting as a mechanism for lowering the danger signal and preserving cells from chronic inflammation. By the way, expression of APOBEC3G was previously shown to be increased by IFNα^80,81^. As shown in **Figure-6F**, we demonstrated that this regulatory mechanism exists also in cardiomyocytes. Interestingly, our 3D-PCR analysis of mtDNA editing revealed a higher rate of mtDNA mutations potentially related to APOBEC3G in E439K-CMs, which is an indirect indication for a more important release of mtDNA into the cytoplasm of mutant cardiomyocytes and/or a higher APOBEC3G activity. Moreover, the expression of the set of genes involved in IFNα/β signaling is increased in E439K-CMs.

Even though the causality between desmin network alterations, sarcomere abnormalities, mitochondrial dysfunctions and/or metabolic perturbations are not well established, it is clear that damages in both desmin and mitochondrial networks precede left ventricular dysfunctions^82^. Our results clearly indicate that a primary effect of desmin E439K mutation in human cardiomyocytes is the mitochondrial abnormalities, which may contribute to the vicious circle leading to progressive dysfunction of cardiomyocytes. This hypothesis is further strengthened by several studies in cell and/or animal models. A comparative study of six pathogenic desmin mutations (S12F, A213V, L345P, A357P, L370P and D399Y) highlighted the role of mitochondrial dysfunction in the pathogenesis of desmin myopathies, independently from the position of the mutation on desmin^15^. Moreover, mitochondrial alterations seem to arise prior to any other observable defect such as loss of contractile material or myofibrillar alignment, according to a study of cardiac tissue from desmin knockout mice^11^. Therefore, these data and our results strongly suggest that mitochondrial dysfunction could be a general mechanism to explain the pathogenesis of desmin-related cardiomyopathies.

It should be noted that mitochondrial dysfunction has been found in several cardiac diseases, *i.e.* dilated cardiomyopathy^83^, hypertrophic cardiomyopathy^84^, arrhythmogenic cardiomyopathy^85^, viral myocarditis^86^, ischemic heart disease^87^ or rare cases of restrictive cardiomyopathy^88^. This clearly suggests that defects in mitochondrial functions and/or structure may be a general trait of cardiac diseases. In that sense, treatments targeting mitochondria appear to be a potential novel approach to strengthen the treatment of this group of conditions as reviewed in^89^. Despite the development of pharmacological molecules targeting key component of mitochondria (reviewed in^90,91^), such as Elamipretide^92^, KL1333^93^ and Idebenone^94^, none of these drugs has yet been approved so far for the treatment of cardiac diseases. Novel research strategies which could lead to the development of innovative treatments are therefore strongly needed. One of the most exciting findings comes from our studies on the transfer of exogenous mitochondria into mutated cardiomyocytes. These experiments reveal that the delivery of functional healthy cardiac mitochondria alleviates the diseased phenotype of desmin-mutated cardiomyocytes. Similar therapeutic approaches have been envisioned to mitigate tissue injury in several cardiac diseases exhibiting dramatic mitochondrial dysfunctions, including ischemic heart diseases in mouse, rat, rabbit and pig models as well as in clinical trials with patients^95–101^. In these settings, exogenous mitochondria have been shown to be internalized by stressed cardiomyocytes and to improve energy metabolism, viability and function of cardiomyocytes^96^. However, despite these positive findings, numerous drawbacks pave the way for the application of mitochondrial transplantation as a regular treatment. In particular, it is difficult to control the integrity and functions of mitochondria in the extra-cellular space or the rate of delivery into cardiomyocytes. Consistent with these concerns, we found that the supply of exogenous functional mitochondria needs to be repeated along the differentiation process of cardiomyocytes to reverse the pathological phenotype indicating that the beneficial effects of these mitochondria are transient. These observations strongly suggest that the transferred mitochondria fail to integrate in the endogenous mitochondrial network of mutant cardiomyocytes but rather they become rapidly defective and probably degraded in the mutant cardiomyocytes.

More recently, beneficial therapeutic effects have been observed in mouse infarcted myocardium following the delivery of mitochondria embedded in microvesicles released by cardiomyocytes derived from iPSCs^62^. The advantage of this approach is to allow a greater functional preservation of the mitochondria that have to be transferred to suffering cardiomyocytes. We applied this method in our study and demonstrated that such an approach is effective in alleviating the genetic defects of desmin-mutated cardiomyocytes as well. Our study raises the question of whether therapeutic efficacy can be influenced by the cellular origin of mitochondria to be transplanted. Consistent with this, only mitochondria isolated from cardiomyocytes have been found to restore contractile functions in a Duchenne muscular dystrophy-driven iPSC-CMs model^102^. This approach appears to be very challenging even if it could be an outstanding opportunity for patients with rare disease where mitochondria is affected. Whatever the fate of the exogenous mitochondria following their internalization in mutated cardiomyocytes, their metabolic effects and their consequences on cardiac function support the conclusion that mitochondrial defects observed in the heart of MFM1 patients critically contribute to the severe symptoms affecting cardiomyocytes. Moreover, our study emphasizes that mitochondria can be considered as a promising therapeutic tool to treat MFM1 myopathy not solely in cardiac muscle but also in all the organs affected by desmin mutations.

## Acknowledgments

We acknowledge Alexandre Simon, Coline Rogue, Gaëlle Revet, Dorota Jeziorowska from CARTHER team for their help in the daily cell culture; Audrey Geeverding and Michaël Trichet from the IBPS electron microscopy core facility and all personnel from the IBPS photon microscopy core facility for helpful advice and technical assistance during microscopy preparation, image acquisition and analysis. The authors thank Brigitte Onteniente and Phenocell SAS (Grasse, France) to generate hiPSCs from MFM1 patient. The authors would also like to express their gratitude to Susanne Bolte from the IBPS photon microscopy core facility, who has sadly died recently.

## Author contributions

Y.H., P.J. and O.A. designed and supervised research; Y.H., Z.L., D.C., R.S., V.B., A.C., J.B., C.L., N.E-J, J-P.V, P.J. and O.A. performed the experiments; Y.H., Z.L., D.C., H.H., A.L., E.K., P.F., J-P.C., G.T., A-M.R., J-P.V., P.J. and O.A. analyzed data; Y.H., P.J. and O.A. wrote the manuscript; all authors proofread the manuscript and approved the final version of this paper.

## Abbreviations

ATP5A: ATP synthase subunit alpha
COX: cytochrome c oxidase
CMs: Cardiomyocytes
Control-CMs: Control cardiomyocytes
Cx43: connexin-43
DCM: dilated cardiomyopathy
DEGs: differentially expressed genes
E439K-CMs: mutated cardiomyocytes
ECAR: extracellular acidification rate
EVs: extracellular vesicles
FCCP: Carbonyl cyanide-4 (trifluoromethoxy) phenylhydrazone
FDR: false discovery rate
FE-SEM: Field emission scanning electron microscopy
GSEA: gene set enrichment analysis
IF: intermediate filament
IFNα/β: Interferon alpha/beta
iPSC: induced pluripotent stem cells
iPSC-CM: cardiomyocytes derived from induced pluripotent stem cells
Iso-E439K-CMs: isogenic mutated cardiomyocytes
MFM: myofibrillar myopathy
MFM1: desmin-related myofibrillar myopathy
MT-COI: mitochondrial cytochrome c oxidase subunit I
mtDNA: mitochondrial DNA
NDUFB8: complex I NADH dehydrogenase (ubiquinone) 1 beta subcomplex subunit 8
OCR: oxygen consumption rate
PBMC: peripheral blood mononucleated cells
PCA: principal component analysis
PFA: paraformaldehyde
SEM: scanning electron microscopy
SDHB: complex II iron-sulfur protein subunit of succinate dehydrogenase
Td: denaturation temperature
TEM: transmission electron microscopy
VDAC: voltage-dependent anion channel
WGCNA: Weighted Gene Correlation Network Analysis

