## Supplemental Tables and Figures for "Critical contribution of mitochondria in the development of cardiomyopathy linked to desmin mutation"

**Short title:** Mitochondria in desmin cardiomyopathy

***Corresponding Authors:** Prof. Onnik Agbulut & Dr. Pierre Joanne. Sorbonne Université, Institut de Biologie Paris-Seine, UMR CNRS 8256, Inserm ERL U1164, 7, quai St Bernard (case 256), F-75005 Paris-France. **&**

### Supplemental Material and methods

**Karyotyping**. The absence of chromosomal aberration was verified by analyzing the karyotype of iPSC cells. Thus, standard chromosomal analyses were performed on cultured derived iPSC clones, by standard procedures [G-banding with trypsin using Giemsa (GTG); R-banding after heat denaturation and Giemsa (RHG)].

**Cytometry.** Cells cultured on Matrigel-coated plates were washed with PBS and detached using Accutase® (for pluripotent stem cells) or TrypLE Select 1X (for cardiomyocytes cultured 4 days after detachment). Immunostaining were performed after incubation in fixation (1% formaldehyde, 20 minutes at RT) and permeabilization (0.5% BSA + 0.1% TritonX-100, 10 minutes) buffers. Then they are stained 15 minutes with anti-Nanog-PE, anti-Sox2-APC and anti-Oct3/4-PerCP-Cy5.5 antibodies (1/100, Human Pluripotent Stem Cell Transcription Factor Analysis Kit, BD). Living cardiomyocytes were stained with MitoTracker Deep Red FM diluted in culture medium (500 nM, 20 min at 37 °C, 5% $CO_2$) or with BODIPY staining solution (2 µM in PBS, 15 min at 37 °C, 5% $CO_2$). Cells are then washed with PBS + 0.5% BSA, resuspended in the same buffer and immediately analyzed on a MACSQuant 10 cytometer. After acquisition, data analysis and figures were performed using FlowJo_v_10.8.1.

**Sequencing.** To confirm the presence of the mutation (or its absence in control iPSC lines) the DNA was extracted using the Genelute Mammalian Genomic DNA kit (Sigma). For PCR, 500 ng of cDNA obtained after extraction of genomic DNA was amplified with 1.5 µM of forward (5'-GAACTAGGAGGGATGGGGAATGT-3') and reverse (5'-TAAAGACAGAGACCCTCTGCCA-3') primers, 1 µM of deoxinucleotide triphosphates, 0.2 µM of MgCl2 and 2.5 units of Taq polymerase (One tas DNA polymerase, NEB). Amplification was performed in a thermal cycler according to the following protocol: 95℃ -30s, then 35 cycles (95℃ -30s, 60 C -30s, 72℃ -1 min) and 72℃ -10 min. Once amplified, electrophoresis of the DNA fragments was performed in a 1.5% agarose gel with TAE buffer and an electric field intensity of 100 V. To stain the DNA in the gel, 1:1000 of GelRED was added during agarose gel preparation. The size of the PCR products was determined by comparison with the size marker (GeneRuler 1 Kb, Thermo Scientific). After migration the DNA bands were purified using the NucleoSpin Gel and PCR clean-up kit (Macherey Nagel) following the manufacturer's instructions. Sequencing was performed by Eurofins Genomics using 5'-GCTGAAGGAAAGGTGTTAAAGTC-3' as forward primer and 5'-CAGCAGCAGCATGAAGTGC-3' as reverse primer.

**Microcontact printing.** A silicon master of the lines was obtained by photolithography techniques and PDMS stamps were fabricated by casting 1:10 PDMS onto the silicon master and baking at 60 °C during 2 hours. Gelatin was activated at room temperature for 30 minutes by incubation with 3.6 mg/mL sodium periodate diluted in in PBS-sodium acetate (50 mM) buffer. For inking, the PDMS stamps were incubated with 0.1% activated gelatin at room temperature for one hour. Then, the solution was aspirated, the stamps were dried and carefully placed in close contact to the surface of a glass coverslip overnight to allow the transfer of protein patterns. The next day, the PDMS stamps were gently removed and the coverslip with the patterns was sterilized for one hour under UV. The coverslips are then immediately used for seeding iPSC-CM.

**Mitochondria Isolation and transfer.** Mitochondria were isolated from Control CMs cells using the Mitochondria Isolation Kit (Thermo Scientific, Waltham, MA, USA, Cat#89874) according to the manufacturer's instructions. The quantity of mitochondria was evaluated by measuring the protein concentration of pelleted mitochondria d using Bicinchoninic Acid Kit (Sigma 1003152319). To evaluate the acute effect of mitondria transfer, pelleted mitochondria corresponding to 0.2 mg per 25 000 cells of proteins were added into the culture medium of CMs for 24 h.

**Supplemental Tables**

**Table S1. List of primary antibodies**

| Target | Antibody | Dilution factor | References |
|---|---|---|---|
| Nanog | Mouse monoclonal (IgG1), clone 1E6C4 | 1:50 | Santa CruzBiotechnology ref # sc-293121 AC |
| TRA1-81 | Mouse monoclonal (IgM) | 1:100 | Merck ref #MAB4381 |
| Desmin | Mouse monoclonal (IgG1), clone D33 | 1:200 | Agilent ref # M076029-2 |
| Desmin | Rabbit monoclonal (IgG), clone Y66 | 1:200 | Abcam ref #ab32362 |
| Desmin | Goat Polyclonal (IgG) | 1:200 | R&D Systems ref #AF3844 |
| Alpha-Actinin | Mouse monoclonal (IgG1), clone EA-53 | 1:200 | Merck ref #A7811 |
| Troponin T | Mouse monoclonal (IgG2a), clone 200805 | 1:1000 | R&D Systems ref #MAB1874 |
| COXIV | Rabbit monoclonal (IgG), clone 3E11 | 1:200 | Cell Signaling ref #4850 |
| VDAC1 | Mouse monoclonal (IgG2b), clone 20B12AF2 | 1:200 | Abcam ref #ab14734 |
| Connexin 43 | Rabbit monoclonal (IgG), clone F-7 | 1:500 | Santa CruzBiotechnology ref # sc-271837 |

**Table S2. List of secondary antibodies**

| Conjugate | Antibody | Dilution factor | References |
|---|---|---|---|
| Alexa 488 | Goat-anti-Mouse IgG2a | 1:1000 | Life Technologies ref # A21131 |
| Alexa 488 | goat anti-Mouse IgG (H+L) | 1:1000 | Life Technologies ref # A11029 |
| Alexa 555 | goat anti mouse IgG1 | 1:1000 | Life Technologies ref # A21127 |
| Alexa 555 | Donkey anti-Goat IgG (H+L) | 1:1000 | Life Technologies ref # A21432 |
| Alexa 647 | goat anti rabbit IgG (H+L) | 1:1000 | Life Technologies ref # A21245 |
| Alexa 568 | goat anti rabbit IgG (H+L) | 1:10000 | Life Technologies ref # A11036 |
| Alexa 488 | goat anti rabbit IgG (H+L) | 1:1000 | Life Technologies ref # A11034 |

### Supplemental Figures with Figure Legends

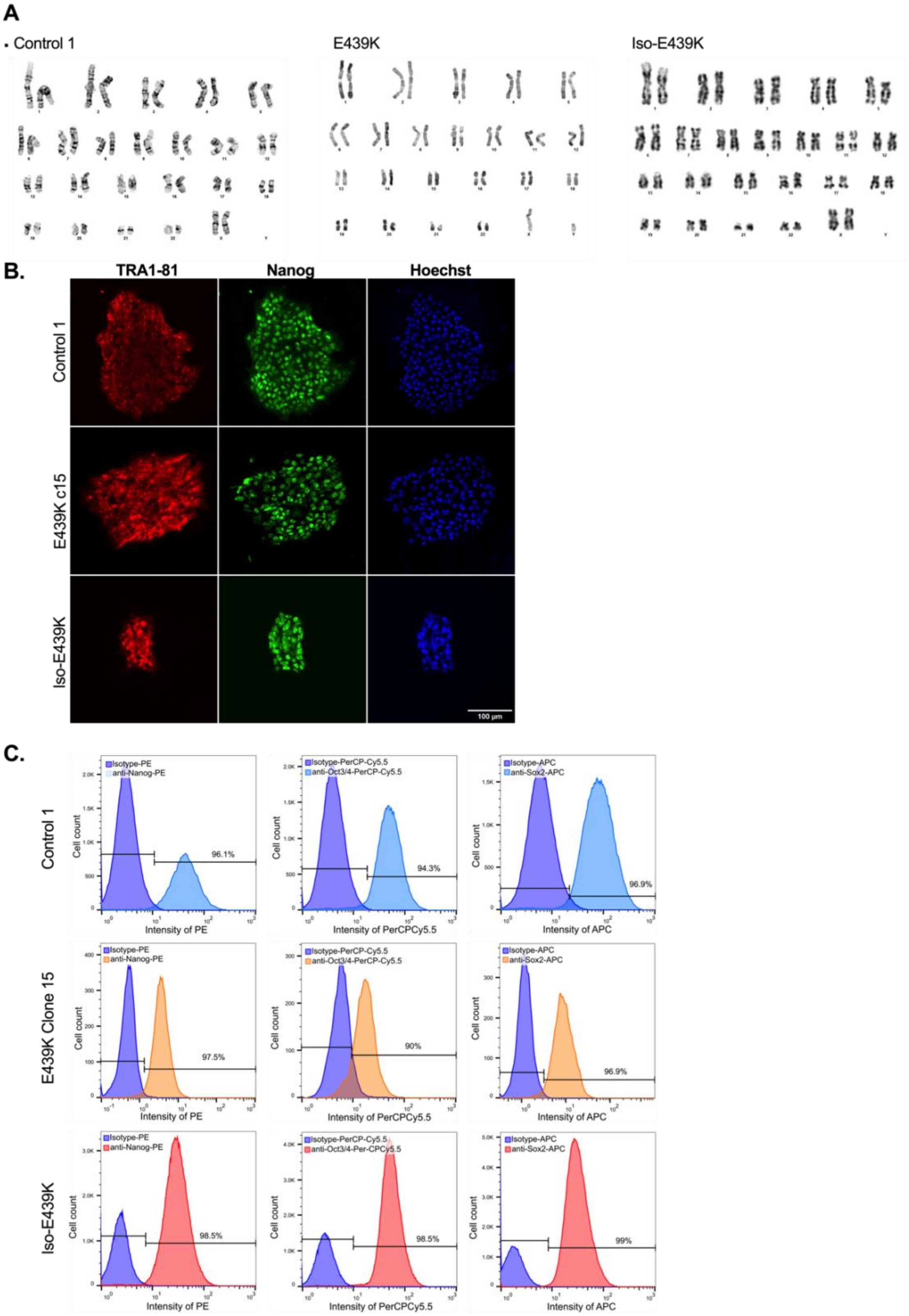


**Figure-S1. Validation of induced pluripotent stem cell lines. (A)** Karyotyping of a control (Control-1), a patient-derived (E439K, clone c15) and the isogenic mutant (Iso-E439K) induced pluripotent stem cell lines. The isogenic mutant was obtained by introducing the point mutation

G-to-A at c.1315 by gene editing using CRISPR-Cas9 (see Material and methods). No genomic abnormalities have been observed in the lines used in this study. **(B)** TRA-1-81 (red) and Nanog (green) immunostaining on growing colonies of a control (Control-1), a patient-derived (E439K, clone c15) and the isogenic mutant (Iso-E439K) induced pluripotent stem cell lines demonstrating correct expression and localization of these pluripotency markers. **(C)** Cytometry analysis of a control (Control-1), a patient-derived (E439K, clone c15) and the isogenic mutant (Iso-E439K) induced pluripotent stem cell lines demonstrating that more than 90% of cells are positive for the pluripotency markers Nanog, Oct3/4 and Sox2.

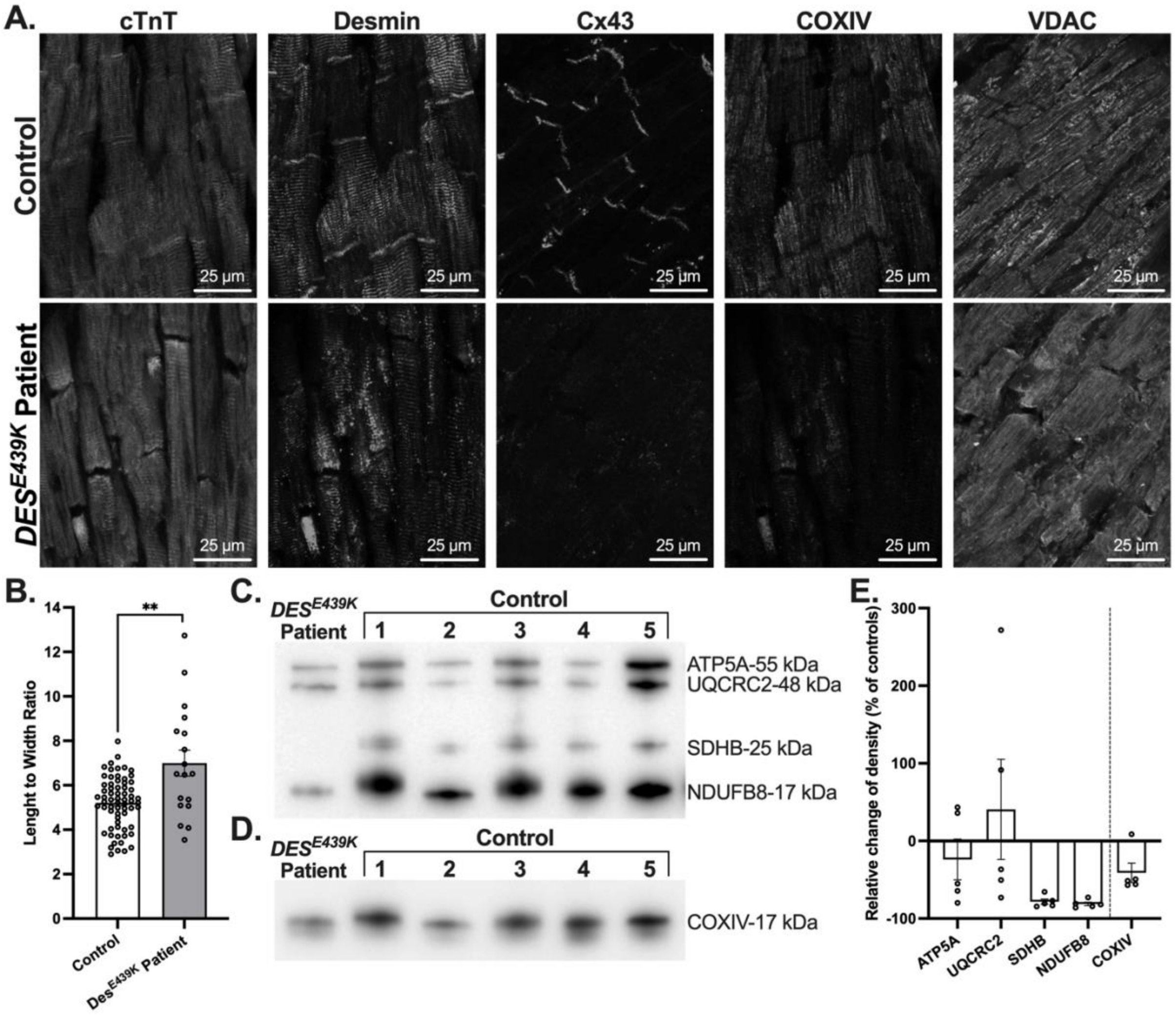


**Figure-S2. Pathological patterns and biochemical analysis of cardiac samples of the Index case CII (see solid arrow in Figure-1A) carrying the *DES*$^{E439K}$ mutation. (A)** Cardiac troponin T (Green), desmin (Red), Connexin 43 (Cx43) (Gray), COXIV (Gray) and VDAC (Gray) immunostaining of formalin-fixed left ventricle sections demonstrating the strong disruption of the desmin network and intercalated discs, and a strong decrease of COXIV and Cx43 labelling in the patient's heart. **(B)** Cardiomyocyte length-to-width ratio was measured using ImageJ software. Values are given as means ± SEM. **, $p < 0.01$. **(C-E)** Western blot analysis (C, D) and densitometry analysis (E) of mitochondrial proteins in control and patient cardiac tissue extracts using a cocktail of antibodies against OXPHOS proteins and COXIV. Note the marked decrease of mitochondrial proteins in the *DES*$^{E439K}$ patient. Results are presented as percent of control density of detected protein bands after normalization to total protein band profile (stain free signal) per lane and are expressed as mean values ± SEM. *, $p < 0.05$, **, $p < 0.01$, ***, $p < 0.001$.

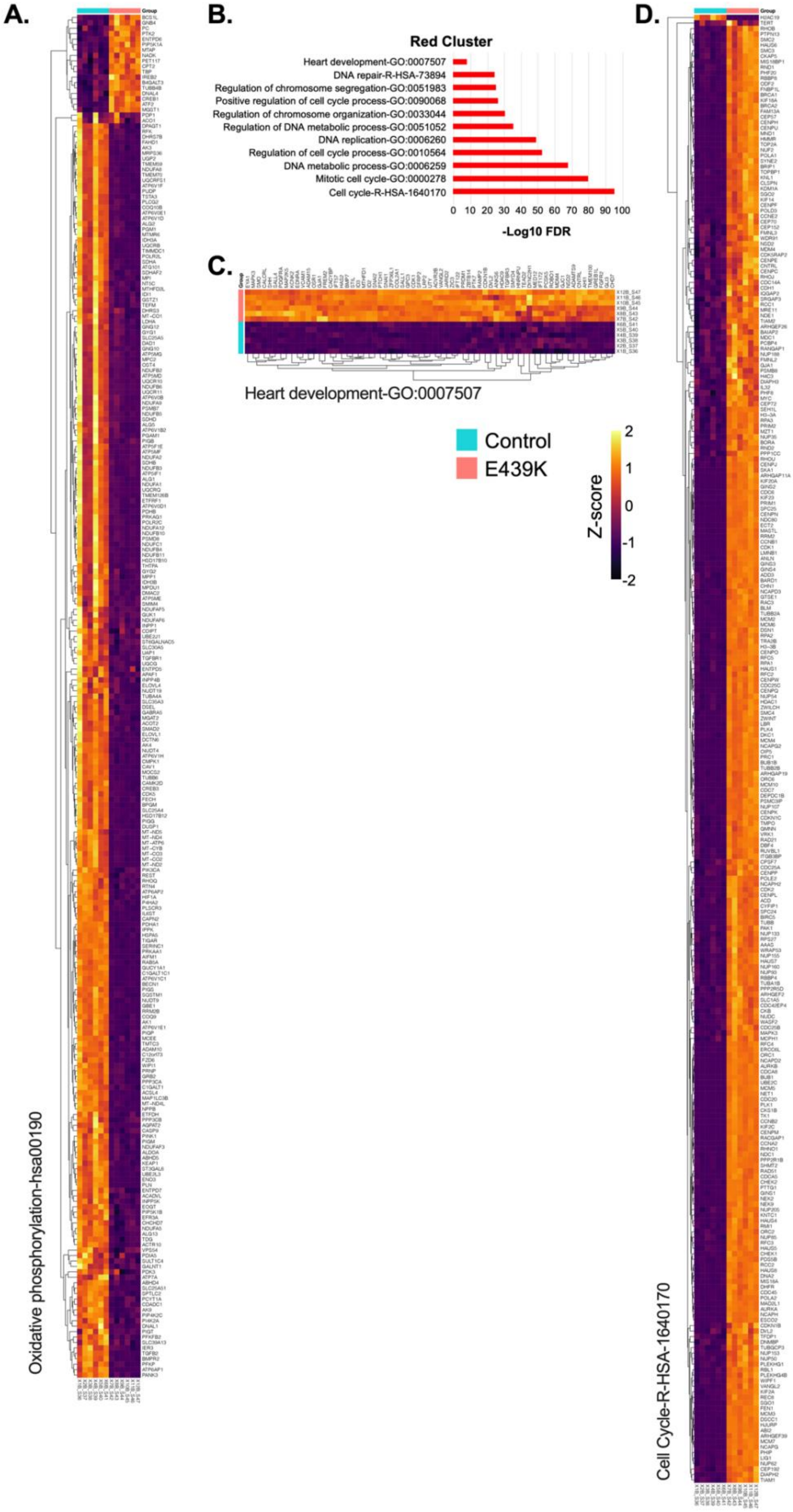
A.
Oxidative phosphorylation-hsa00190
B.
Red Cluster
Heart development-GO:0007507
DNA repair-R-HSA-73894
Regulation of chromosome segregation-GO:0051983
Positive regulation of cell cycle process-GO:0090068
Regulation of chromosome organization-GO:0033044
Regulation of DNA metabolic process-GO:0051052
DNA replication-GO:0006260
Regulation of cell cycle process-GO:0010564
DNA metabolic process-GO:0006259
Mitotic cell cycle-GO:0000278
Cell cycle-R-HSA-1640170
0 10 20 30 40 50 60 70 80 90 100
-Log10 FDR
C.
Heart development-GO:0007507
Control
E439K
Z-score
2
1
0
-1
-2
D.
Cell Cycle-R-HSA-1640170

**Figure-S3. Transcriptomic analysis of E439K-CMs compared to Control-CMs. (A)** Set of gene related to oxidative phosphorylation analyzed after gene set enrichment analysis of the genes of the Blue cluster determine by unsupervised WGCNA. Moreover, the enrichment analysis of the Red cluster **(B)** reveals different sets of genes related to heart development **(C)** and Cell Cycle **(D)**.

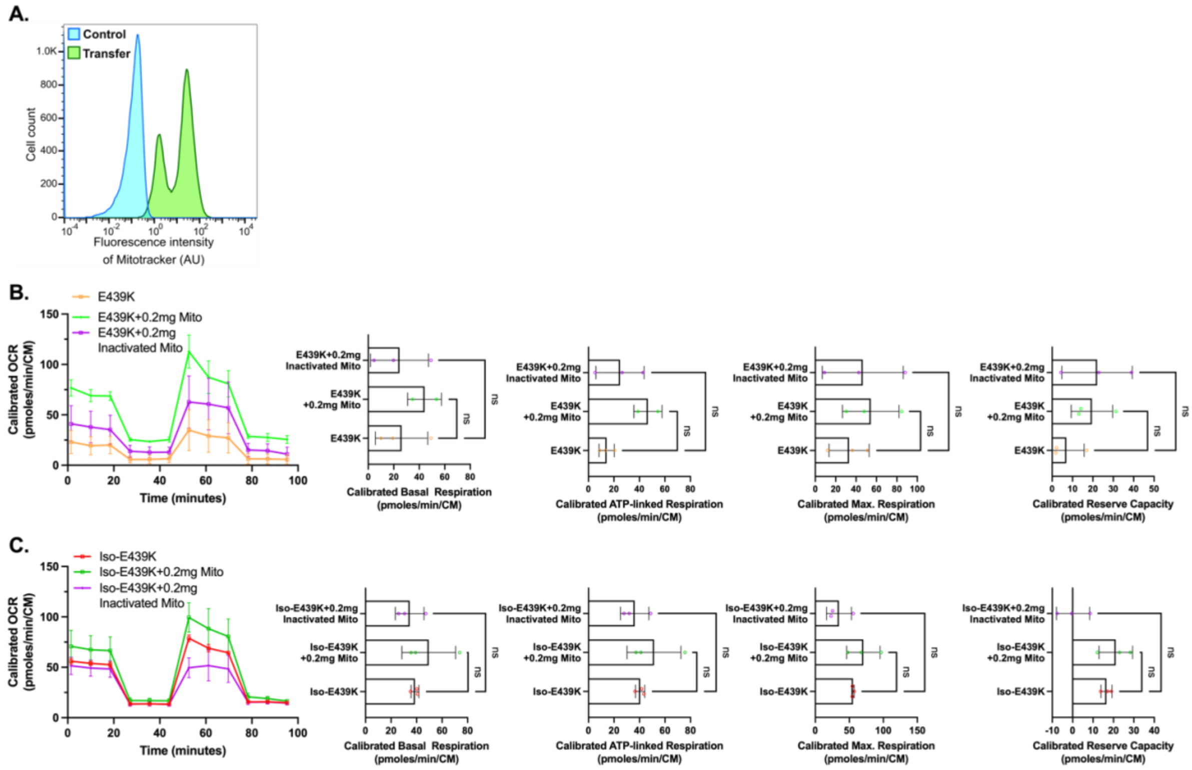


**Figure-S4. Effect of acute treatment of cardiomyocytes with mitochondria purified from Control-CMs. (A)** 24h after the treatment of Control-CMs derived mitochondria stained with MitoTracker®, cytometry analysis reveals that the mitochondria are effectively transferred into cardiomyocytes. **(B-C)** Representative mitochondrial oxygen consumption rate (OCR) profiles in E439K-CMs **(B)** or Iso-E439K-CMs **(C)** 24 hours after the transfer of mitochondria reveal that this treatment can modulate the respiration of mutant cardiomyocytes. OCR profiles are expressed as pmol O2/min normalized to the number of cardiomyocytes and calibrated to the lowest value of the plate after addition of rotenone and antimycine. For each mutant line, quantification of basal respiration, ATP-linked respiration, maximal respiration and reserve capacity were calculated. Values are expressed as mean ± SEM. *, $p < 0.05$, **, $p < 0.01$, ***, $p < 0.001$.

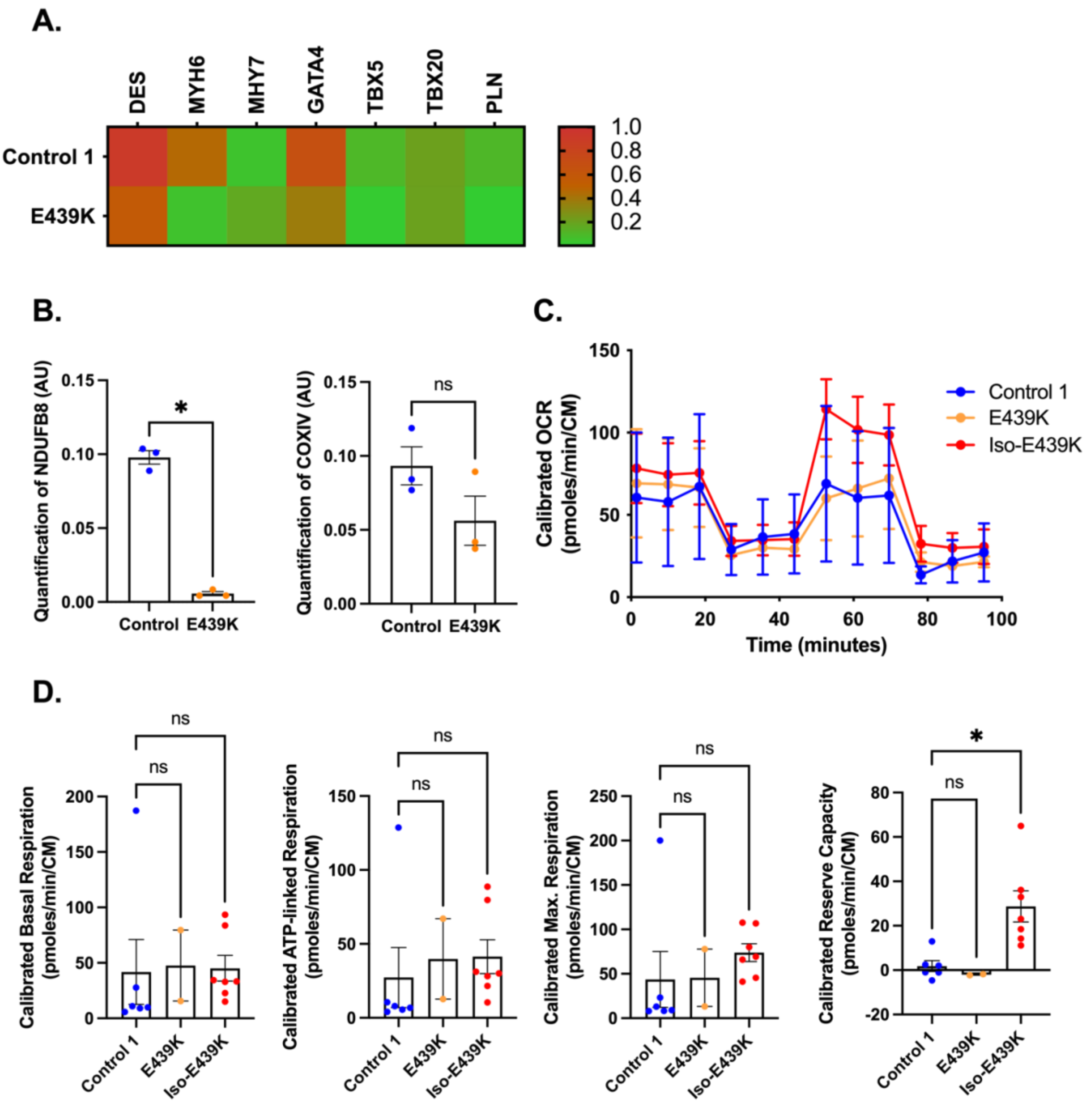


**Figure-S5. Absence of strong differences at the beginning of differentiation in terms of genes and proteins expression as well as respiratory function. (A)** Heatmap op the gene expression 5 days after the initiation of cardiac differentiation of Control 1 and E439K iPSC lines revealing few differences in the expression of cardiomyocytes-related genes. **(B)** Quantitative analysis of Western Blot of NDUFB8 and COXIV showing a decrease of NDUFB8 mitochondrial protein but not COXIV in E439K-CMs compared to Control-CMs 7 days after the initiation of differentiation. Results are presented as fold change to control after normalizing to Stain-Free profile. **(C)** Representative mitochondrial oxygen consumption rate (OCR) profiles in Control-CMs, E439K-CMs and Iso-E439K-CMs 8 days after the initiation of differentiation. OCR profiles are expressed as pmol $O_2$/min normalized to the number of cardiomyocytes and calibrated to the lowest value of the plate after addition of rotenone and antimycin A. **(D)** Quantification of basal respiration, ATP-linked respiration, maximal respiration and reserve capacity of cardiomyocytes. All values are expressed as mean ± SEM. *, $p < 0.05$, **, $p < 0.01$, ***, $p < 0.001$.
